# Heterogeneity in viral replication dynamics shapes the antiviral response

**DOI:** 10.1101/2022.06.08.495262

**Authors:** L.J.M. Bruurs, M. Müller, J.G. Schipper, H.H. Rabouw, S. Boersma, F.J.M. van Kuppeveld, M.E. Tanenbaum

## Abstract

In response to virus infection, host cells can activate antiviral signaling to restrict virus replication and communicate viral infection to neighboring cells. For poorly understood reasons, antiviral response activation is highly heterogeneous among infected cells; both quantitatively (level of pathway activation) and qualitatively (transcribed antiviral gene set). Here, we used live-cell single-molecule imaging to simultaneously visualize viral infection and antiviral signaling, providing quantitative insights into antiviral response activation in single cells; first, the probability of activating an antiviral response varies throughout infection, with most efficient activation occurring several hours after the first viral replication. Second, cell-to-cell heterogeneity in viral replication rates early in infection determine the efficiency of antiviral response activation. Finally, variation in signaling strength of the viral sensing pathway result in qualitatively distinct antiviral responses. Together, this works identifies key parameters that shape the antiviral response and provides quantitative insights into the origin of heterogeneity in the antiviral response.

## Introduction

The innate immune system provides a first line of defense against viral infection and stimulates activation of the adaptive immune system (O’Neill and Bowie, 2010; Schoggins and Rice, 2011). A key step in innate immune activation is the production of type I interferons (IFN), which are important signaling molecules that induce an antiviral state in neighboring cells and thereby protect these cells against viral infection (Mesev et al., 2019; Schoggins and Rice, 2011). However, excessive activation of IFN signaling can be toxic to tissues and can contribute to hyperinflammation, a condition that underlies various pathologies, including respiratory viral diseases like COVID-19 (Lee and Shin, 2020; Nelemans and Kikkert, 2019; Postal et al., 2020; Sposito et al., 2021). Therefore, the ability to precisely tune the IFN-based antiviral response during infection is essential to restrict virus infection while preventing excessive inflammation. Furthermore, in the absence of viral infection, stringent control of antiviral response activation is required to prevent initiation of a spurious response, which can cause, among others, a spectrum of syndromes collectively known as ‘interferonopathies’ (Crow, 2011; Crow and Stetson, 2021).

Initiation of an antiviral response depends on detection of viral infection by the host cell (O’Neill and Bowie, 2010). For RNA viruses, double stranded RNA (dsRNA), which is often formed during viral replication, constitutes an important ligand for activating the cellular antiviral response (Pichlmair et al., 2009). Members of the family of RIG-I-like receptors (RLRs), including melanoma differentiation-associated protein 5 (MDA5), are able to sense cytosolic dsRNA (Dias Junior et al., 2019; Pichlmair et al., 2009; Rehwinkel and Gack, 2020). Binding of MDA5 to dsRNA activates a dsRNA sensing signaling pathway that culminates in nuclear translocation of the interferon regulatory factor (IRF) family of transcription factors (e.g. IRF3 and IRF7) (O’Neill and Bowie, 2010). Nuclear IRFs induce transcription of several genes with antiviral functions (e.g. IFIT1 and viperin) as well as secreted proinflammatory cytokines, including IFNs. Upon binding to the interferon α/β receptor, either via autocrine and paracrine signaling, IFNs induce expression of a largely distinct set of antiviral genes, collectively known as interferon stimulated genes (ISGs), that can effectively protect neighboring cells against virus infection (Andersen et al., 2008; Grandvaux et al., 2002; Savitsky et al., 2010; Schneider et al., 2014; Schoggins, 2019; Schoggins and Rice, 2011). Activation of the dsRNA sensing pathway therefore results in antiviral gene expression in the infected cell and can induce protective ISG expression in uninfected neighboring cells.

To prevent an antiviral response by the host cell, viruses have evolved strategies to repress activation of the IFN pathway and to inhibit expression of antiviral genes (here referred to as viral antagonism) (Feng et al., 2014; Lei et al., 2016). However, this viral antagonism is not always fully effective in preventing activation of the antiviral response (Rand et al., 2014), and a subset of infected cells are generally capable of launching an antiviral response and expressing IRF target genes, resulting in cell-to-cell heterogeneity in the antiviral response. For instance, infections with different viruses result in IFNB1 production in <1% −30% of infected cells (Doğanay et al., 2017; Drayman et al., 2019; Patil et al., 2015; Zawatzky et al., 1985; Zhao et al., 2012). The antiviral response can also differ *qualitatively* among infected cells; the set of transcribed antiviral genes can vary between infected cells of the same cell type (Doğanay et al., 2017; Drayman et al., 2019; Sjaastad et al., 2018), creating an additional layer of cell-to-cell heterogeneity in the antiviral response.

While heterogeneity in the antiviral response has been widely reported and likely plays a major role in controlling viral spread, factors that determine this heterogenous response to viral infection remain poorly understood. Exogenous overexpression of various host cell antiviral proteins (MDA5, RIG-I, IRF7) was found to increase the fraction of IFNB1 producing cells (Doğanay et al., 2017; Zhao et al., 2012), suggesting that expression levels of these proteins may affect the efficiency of antiviral response activation. However, it is unclear whether *endogenous* expression levels of these factors vary between cells, and whether the extent of such variation is sufficient to explain the observed heterogeneity in antiviral response activation (Wimmers et al., 2018). In fact, cell-to-cell heterogeneity in the antiviral response has even been reported in sister cells after cell division which likely have very similar gene expression levels, suggesting that factors other than host gene expression differences may be important in causing heterogeneity in the antiviral response (Rand et al., 2012; Talemi and Höfer, 2018).

As for the antiviral response, considerable variation also exists in the progression of viral infection among infected cells. For example, viral replication rates vary strongly among infected cells, possibly as a result of differences in the infecting virus (e.g. genetic variation in viral genome sequence), and through differences in the host cell (e.g. expression levels of restriction factors) (Guo et al., 2017; Jones et al., 2021; Russell et al., 2018; Schulte and Andino, 2014). Since both viral replication rates and antiviral response show substantial cell-to-cell heterogeneity, one possibility is that heterogeneity in viral replication is causally linked to heterogeneity in antiviral signaling. However, the conclusions of studies assessing this correlation are conflicting, with some studies finding a positive correlation between viral load and antiviral signaling (Fiege et al., 2021; Zhao et al., 2012), while others finding a negative (Drayman et al., 2019; O’Neal et al., 2019) or no correlation (Doğanay et al., 2017; Patil et al., 2015). Importantly, most of these studies make use of single time point ‘snapshot’ measurements (e.g. qPCR, FISH or RNA-seq) to determine viral genome abundance. A major limitation of such measurements is that variability in the start of infection in different cells cannot be taken into account. Assessing the moment of infection is critical to discriminate cells with low viral replication rates from cells in which infection initiated later in the experiment. Moreover, several studies used infections with an MOI>1 (Doğanay et al., 2017; Zawatzky et al., 1985; Zhao et al., 2012), resulting in considerable variation in the number of virions that infect a single cell. Because MOI affects the rate of virus replication (Martin et al., 2020; Schulte and Andino, 2014), it is challenging to disentangle heterogeneity in viral replication rates from variation in the number of infecting virions in infections with MOI>1. Thus, to study the effect of viral replication rates on innate immune activation, highly sensitive live-cell read-outs are required to precisely determine the moment of infection by individual viruses and the timing and strength of antiviral response activation in single cells.

Here, we combine <u>V</u>irus Infection <u>R</u>eal-time <u>IM</u>aging (VIRIM), a live-cell single-molecule imaging method that we recently developed for detecting viral infection and replication in living cells (Boersma et al., 2020), with real-time, highly sensitive read-outs of the dsRNA sensing pathway and antiviral response activation. Using encephalomyocarditis virus (EMCV), a member of the picornavirus family, we uncover multiple parameters that contribute to cell-to-cell heterogeneity in the antiviral response. First, we show that the probability of antiviral response activation varies during infection and is highest several hours after the first round of viral replication. Second, we find that antiviral response activation is more likely to occur in cells that exhibit low viral replication rates during the first hours of infection. Finally, we identify qualitative differences in the antiviral response (i.e. expression of different antiviral genes) among infected cells and show that these differences are correlated with variation in the signaling strength of the dsRNA sensing pathway. Thus, through highly sensitive real-time imaging this work uncovers how variation in infection and host cell sensing kinetics shape the antiviral response.

## Results

### Real-time imaging of EMCV infection

To generate a quantitative framework for understanding viral replication and antiviral response kinetics, we made use of the (+)ssRNA encephalomyocarditis virus (EMCV). This member of the picornavirus family has been used extensively to study antiviral response mechanisms (Deddouche et al., 2014; Feng et al., 2014; Fout and Simon, 1983; Hato et al., 2007; Satoh et al., 2010) and we previously demonstrated that it is amenable to the VIRIM assay (Boersma et al., 2020). In brief, the VIRIM assay uses two components to visualize virus infection; first, 5 copies of a short peptide called the SunTag (Tanenbaum et al., 2014) are inserted at the N-terminus of the viral polyprotein (5xSunTag-EMCV). C-terminal to the 5xSunTag array, a cleavage site is introduced for the viral 3C(D) protease to enable release of the 5xSunTag array from the mature viral polyprotein in order to prevent the SunTag array from interfering with viral protein functions. The second component of VIRIM consists of a genetically-encoded single chain variable fragment (scFv) antibody that binds tightly to the SunTag peptide (referred to as <u>S</u>un<u>T</u>ag <u>a</u>nti<u>b</u>ody, STAb) and that is fused to a fluorescent protein (GFP-STAb). When the engineered 5xSunTag-EMCV infects cells that express GFP-STAb and the viral genome is translated in the host cell cytoplasm, SunTag peptides are produced and are co-translationally bound by the GFP-STAb (Fig. 1A). As each ribosome translating the 5xSunTag-EMCV genome is associated with 5 SunTag peptides (and thus 5 STAb-GFPs), and because multiple ribosomes translate the 5xSunTag-EMCV genome simultaneously, many GFP-STAb molecules are recruited to a translating viral RNA (vRNA), resulting in a bright fluorescent spot that can be detected by spinning disk confocal microscopy. Single ‘mature’ (i.e. ribosome released) SunTag polypeptides are not detectable as fluorescent foci, due to their much weaker fluorescence (Boersma et al., 2020). Importantly, translation of the incoming viral genome occurs very rapidly, within minutes after entry into the cytoplasm, and the majority of genomes undergo translation (Boersma et al., 2020). Thus, the number of fluorescent SunTag GFP foci in a cell accurately reports on the number of viral genomes and can therefore be used both to determine the start of viral translation and to assess viral replication kinetics early in infection (Boersma et al., 2020).

**Fig 1.**
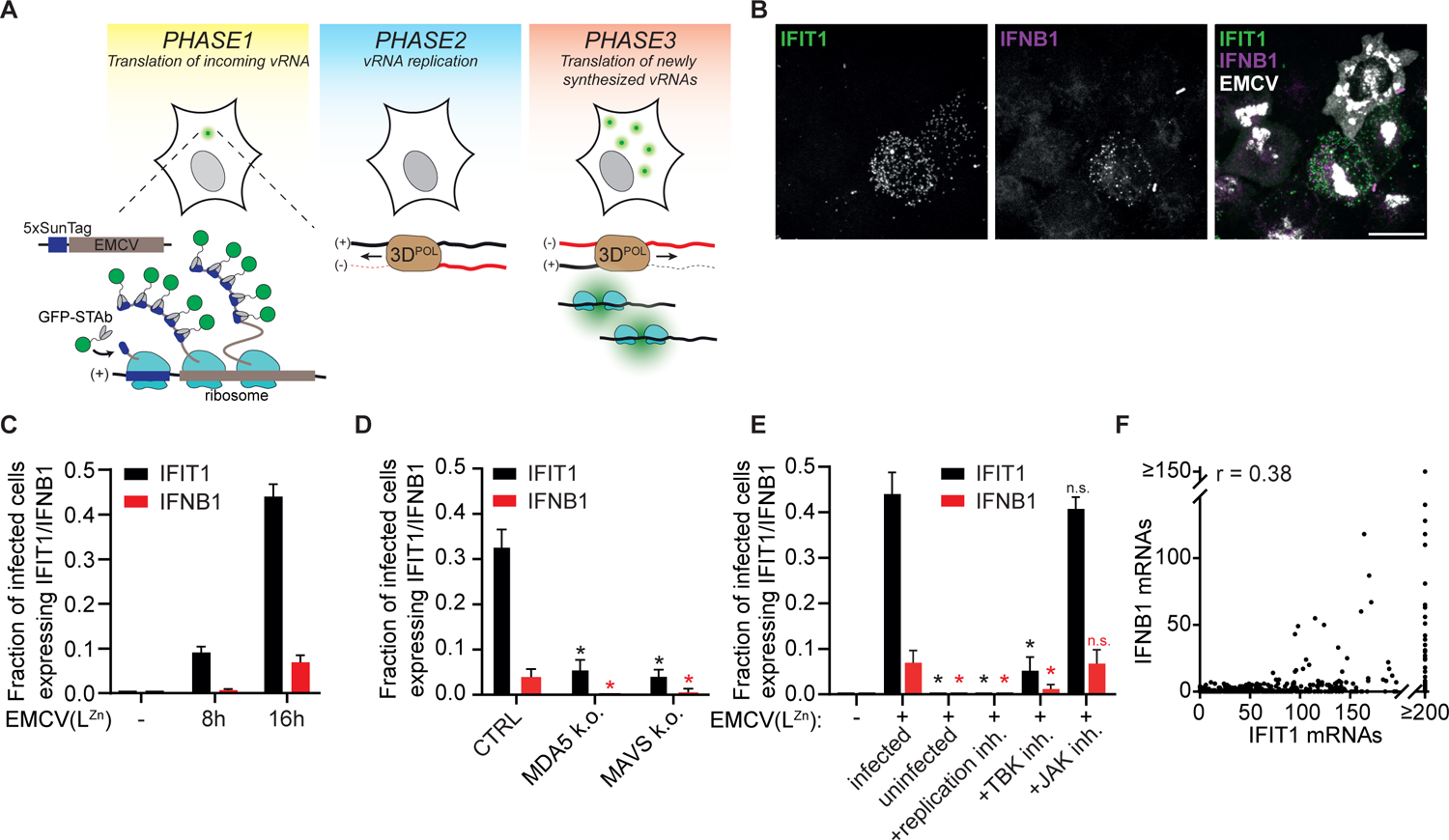
IFIT1 is expressed in a subset of EMCV(L^Zn^) infected cells in an MDA5/MAVS/TBK dependent manner. A) Schematic representation of VIRIM experimental setup and VIRIM phases. During phase 1, a single GFP spot is visible, which represents the incoming vRNA that is translated by the host cell translation machinery. In phase 2 translation of the incoming genome is shut down and the vRNA undergoes replication, resulting in the disappearance of the GFP spot. In phase 3, newly-synthesized daughter vRNAs are produced, which are rapidly initiate translation, resulting in the appearance of multiple new GFP spots. 3D^POL^ = EMCV RNA-dependent RNA polymerase B) Representative smFISH image of 5xSunTag-EMCV(L^Zn^) infected HeLa cells at 16 h.p.i. labeled with probes targeting IFIT1 and IFNB1 mRNAs and viral EMCV genomes. Scale bar indicates 20 µm. C) Fraction of infected cells with more than 10 IFNB1 (red bars) or 20 IFIT1 (black bars) mRNAs at 0, 8 and 16 h.p.i. (n = 3 independent experiments). D) Fraction of infected cells expressing more than 10 IFNB1 or 20 IFIT1 mRNAs at 16 h.p.i. in either HeLa control or MDA5 and MAVS k.o. HeLa cells (n = 4 independent experiments). * indicates p < 0.05 using independent samples T-test compared to HeLa CTRL. Black * or n.s. indicate condition was compared with the fraction of IFIT1+ cells in the infected condition, red * or n.s. was compared with the fraction IFNB1+ cells. E) Fraction of infected cells expressing more than 10 IFNB1 or 20 IFIT1 mRNAs at 16 h.p.i. with or without treatment with an EMCV replication inhibitor (dipyridamole, DiP), TBK1/IKKε inhibitor (MRT67307, MRT.) or JAK1/3 inhibitor (Tofacitinib, TOFA.) (n = 3 independent experiments). * indicates p < 0.05 using independent samples T-test compared to EMCV(L^Zn^) infected, n.s. indicates non significant F) Scatter plot showing the number of IFIT1 and IFNB1 mRNAs in 5xSunTag-EMCV(L^Zn^) infected cells 16 h.p.i. (r indicates Pearson’s correlation coefficient, n = 641 cells from 6 independent experiments). Bars and error bars indicate average ± s.e.m. in all panels

EMCV infection leads to potent inhibition of the dsRNA sensing pathway, thereby preventing expression of IRF3 target genes (Hato et al., 2007), and limiting our ability to study viral sensing and activation of the antiviral response pathway. Although several EMCV proteins are implicated in suppressing the dsRNA sensing pathway, the N-terminal Leader (L) protein is considered the main IFN antagonist of EMCV (Han et al., 2021; Hato et al., 2007; Huang et al., 2017; Li et al., 2019). Indeed, an EMCV virus with inactivating mutations in the zinc finger domain of the L protein (“EMCV(L^Zn^)”) induces potent expression of antiviral genes (Hato et al., 2007), which we could confirm by smFISH (Fig. S1A). Importantly, EMCV(L^Zn^) infection induced antiviral gene expression only in a subset of infected cells, indicating that L protein inactivation does not result in a loss of cell-to-cell heterogeneity in antiviral response activation (Fig. 1B, S1A). Furthermore, introduction of the 5xSunTag array into the viral genome did not affect the efficiency of antiviral response activation (Fig. S1B). Performing VIRIM using 5xSunTag-EMCV(L^Zn^) virus (validated by analysis of EMCV(L^WT^)) therefore constitutes a powerful approach to study heterogeneity in the antiviral response.

### Single cell analysis of IFIT1 and IFNB1 expression reveals heterogeneous antiviral responses

To sensitively monitor activation of the antiviral response, we searched for genes that are transcriptionally activated in cells that have sensed viral dsRNA via the MDA5/MAVS/IRF pathway. IFNB1, the best known IRF3 target gene, is expressed only in a subset of cells in which IRF3 is activated (Doğanay et al., 2017) and is thus not an ideal marker gene to report on antiviral signaling activation. Therefore, we set out to identify a different marker gene that better reports on cells in which the dsRNA sensing pathway is activated. We focused on the gene interferon induced protein with tetratricopeptide repeats 1 (IFIT1). While IFIT1 is best known as an ISG, IFIT1 expression is also upregulated by IRF3-dependent transcription in an IFN-independent manner (Bandyopadhyay et al., 1995; Diamond and Farzan, 2013; Grandvaux et al., 2002). IFIT1 transcription is strongly induced during virus infection and induction of IFIT1 expression can be detected well before IFNB1 expression (Doğanay et al., 2017).

We assessed IFIT1 and IFNB1 expression in HeLa cells in response to 5xSunTag-EMCV(L^Zn^) infection using single molecule fluorescence in situ hybridization (smFISH), a highly sensitive, single-cell method for analysis of gene expression. smFISH probes targeting IFIT1 and IFNB1 mRNAs were combined with probes targeting the EMCV genome to identify infected cells (Fig. 1B). Baseline expression of both IFIT1 and IFNB1 in uninfected HeLa cells is very low (98% of uninfected cells have <4 IFIT1 and <2 IFNB1 mRNAs) (Fig. S1C). Based on the number of smFISH spots in uninfected cells we set a stringent cut-off value of 20 IFIT1 and 10 IFNB1 mRNAs, above which a cell is called positive for expression of either gene. Both IFIT1 and IFNB1 expression were strongly induced by viral infection, and we found that the fraction of cells expressing IFIT1 was substantially higher than the fraction expressing IFNB1 at multiple time points in infection (9.1% and 0.7% at 8 h.p.i. and 44.0% and 6.9% at 16 h.p.i. for IFIT1 and IFNB1, respectively) (Fig. 1C). This indicates that analysis of IFIT1 expression indeed provides a more sensitive readout of antiviral signaling activation. Notably, even at the latest time point (16 h.p.i.), after which we observed extensive cell death of EMCV-infected cells, less than half of the infected cells showed IFIT1 expression, indicating that antiviral response activation following 5xSunTag-EMCV(L^Zn^) infection is indeed heterogeneous.

Through multiple lines of investigation, we confirmed that expression of IFIT1 (and IFNB1) is a direct consequence of detection of viral dsRNA in the infected host cell rather than of paracrine IFN signaling: first, deletion of the dsRNA sensor MDA5 or downstream inactivation of the dsRNA sensing pathway, by deletion of MAVS or inhibition of TBK1 (the kinase responsible for IRF3 activation), resulted in a strong reduction of IFIT1 and IFNB1 expression (Fig. 1D,E). In contrast, inhibition of paracrine IFN signaling through JAK inhibition did not affect expression of IFIT1 and IFNB1 in response to EMCV infection (Fig. 1E, S1D). Second, IFIT1 and IFNB1 expression required viral replication, as inhibition of EMCV replication by dipyridamole (DiP) (Fata-Hartley and Palmenberg, 2005) results in complete loss of their expression (Fig. 1E). Lastly, expression of IFIT1 is limited to infected cells and is not observed in neighboring uninfected cells (Fig. 1E,”uninfected”). Together, these experiments establish IFIT1 expression as a sensitive marker for cells that sense intracellular infection through viral dsRNA and activate an antiviral response.

Simultaneous smFISH labeling of IFIT1 and IFNB1 mRNAs in single cells revealed that all IFNB1+ cells also express IFIT1, but not all IFIT1+ cells express IFNB1, confirming that IFNB1+ cells constitute a subset of IFIT1+ cells (Fig. 1F). The absence of IFNB1 mRNAs in a subset of IFIT1+ cells is unlikely the result of poor smFISH labeling efficiency in IFIT1+/IFNB1-cells, as the IFIT1 smFISH spot intensity is similar in IFNB1- and IFNB1+ cells (Fig. S1E). Interestingly, IFNB1 expression is mostly observed in cells expressing high levels of IFIT1 (Fig. 1F), suggesting that only cells with very strong antiviral response activation preferentially induce IFNB1 transcription. Combining smFISH for IFIT1 and IFNB1 therefore reveals heterogeneity in host cell responses and allows identification of at least three quantitatively and qualitatively distinct host responses to viral infection: 1) no antiviral response activation (IFIT1-/IFNB1-), 2) activation of IFIT1 expression only (IFIT1+/IFNB1-) and 3) activation of both IFIT1 and IFNB1 expression (IFIT1+/IFNB1+).

### Cells that activate antiviral gene expression have lower viral loads

Having established a smFISH-based approach that allows visualization of heterogeneous antiviral responses, we next combined this smFISH-based approach with live-cell VIRIM (by performing VIRIM first, followed by smFISH analysis of the same cells) in order to assess whether variation in viral replication dynamics can explain heterogeneity in antiviral response. Based on VIRIM, we previously categorized early infection into several distinct ‘phases’ of infection (See fig.1A) (Boersma et al., 2020); upon entry of the ‘mother’ vRNA into the host cell cytoplasm, the vRNA rapidly undergoes translation, which is observable as a single fluorescent spot (referred to as “phase 1”). Phase 1 is followed by vRNA translation shut down and initiation of vRNA replication, at which time the GFP spot disappears (“phase 2”). The subsequent synthesis of multiple daughter vRNAs that rapidly initiate translation is observed as a rapid, sequential appearance of multiple GFP foci (“phase 3”). Thus, the first vRNA replication initiates in phase 2, and the appearance of multiple GFP foci in phase 3 indicates that viral replication has occurred. Therefore, the start of phase 3 reports on the presence of the first molecule of viral dsRNA in the cell and thus marks the first opportunity for viral sensing by the host cell.

We combined live-cell EMCV VIRIM with smFISH for IFIT1 and IFNB1 mRNAs and EMCV genomes; infections were imaged for 16 h using VIRIM, after which cells were fixed and smFISH staining and analysis was performed on the same cells as imaged using VIRIM. This combined approach enabled compiling of live infection data, antiviral gene expression and late-stage viral load of individual infected cells (Fig. 2A, Video S1). Because not all infections initiate simultaneously, the duration of infection at the moment of cell fixation is highly variable among infected cells (Fig. S2A). To correct for differences in infection duration in individual cells, all infections were aligned *in silico* to the start of VIRIM phase 3, which approximates the first moment in infection when viral dsRNA is present in the cell. Synchronizing all infections to the start of phase 3 revealed that a considerable lag period exists between the first round of virus replication and the emergence of IFIT1 and IFNB1 transcripts (a lag of approximately 7 h) (Fig. 2B-D). The timing of IFIT1 and IFNB1 mRNA expression was confirmed by qPCR at varying time points during infection (Fig. S2B). Of note, whereas IFIT1 induction is first detected at 8 - 10 h.p.i. by qPCR, it was only detected 7 h after the start of phase 3 by smFISH (Fig. 2D). On average infections require 6 h to complete phase 3 after addition of virus (Fig. S2A) and the induction as observed by smFISH would thus appear to occur considerably later than as detected by qPCR. However, a subset of infections rapidly complete the initial replication (<2,5 h) and the moment of antiviral gene induction as detected by qPCR likely reflects activation of IFIT1 expression in this subset of cells. Together, these findings reveal a substantial time delay between completion of the initial vRNA replication and the expression of antiviral genes.

**Fig 2.**
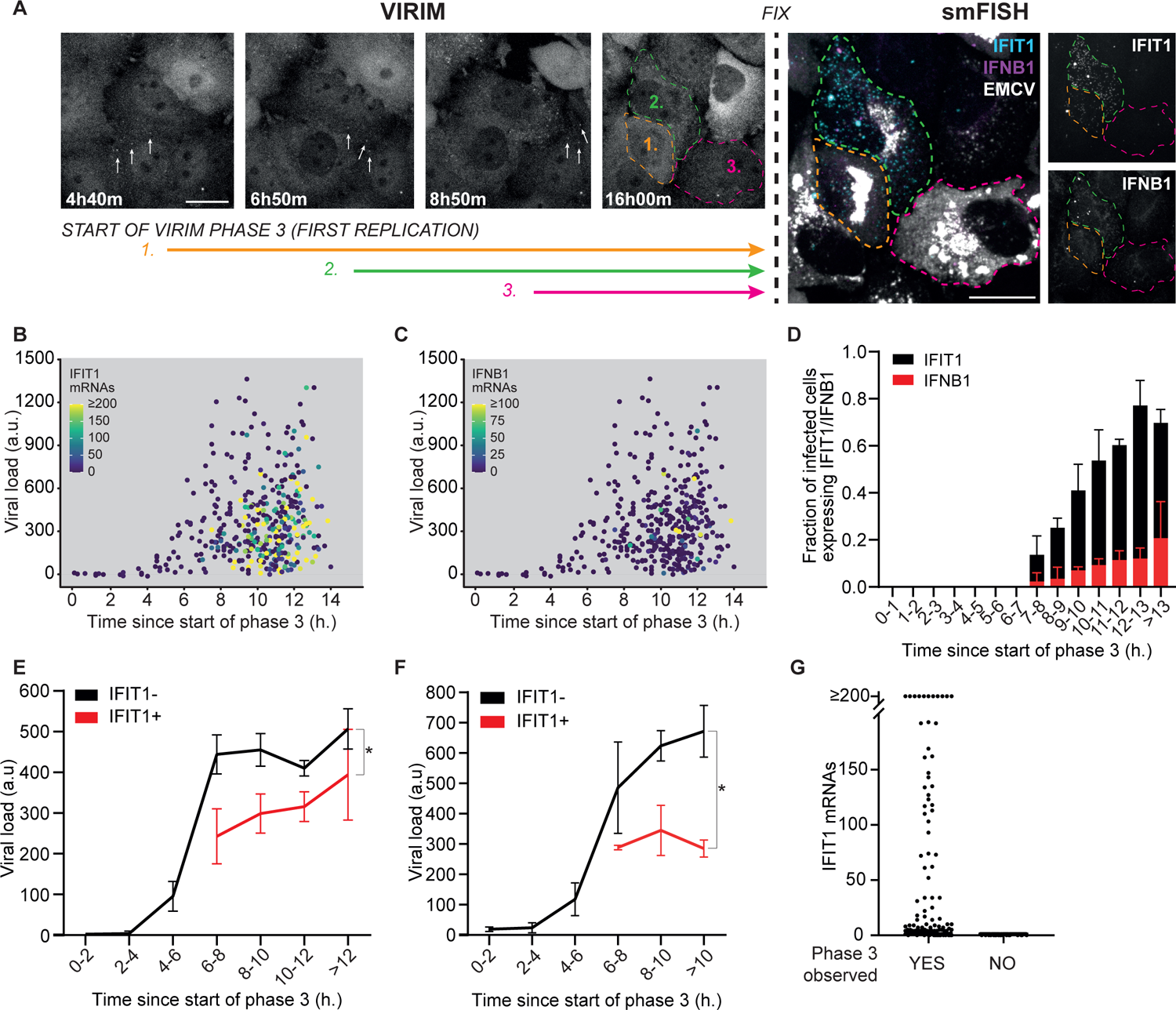
An antiviral response is preferentially activated in cells that have a lower viral load. For all panels, HeLa cells expressing STAb-GFP were infected with 5xSunTag-EMCV(L^Zn^) and imaged for 16 h. After that cells were fixed and subjected to smFISH labeling using probes targeting IFIT1 and IFNB1 mRNAs or EMCV genomes. A) Representative images of live-cell virus imaging using VIRIM combined with smFISH for IFIT1, IFNB1 and EMCV. Left (VIRIM): Images from a movie of three infected cells in which the first vRNA replication occurs at different time points after virus addition. Time indicates time since virus addition. White arrows highlight viral translation sites. Colored arrows indicate the period during which dsRNA is present in the infected cell. Right (smFISH): smFISH labeling of the three infected cells with probes targeting IFNB1 and IFIT1 mRNA and EMCV genomes. Colored dotted lines mark the outline of cell. Scale bar indicates 20 µm. B) Scatter plot showing viral load relative to the time in infection (relative to the start of phase 3). Spot color indicates number of IFIT1 mRNAs. (n = 399 cells in 3 independent experiments) C) Scatter plot showing viral load relative to the time in infection (relative to the start of phase 3). Spot color indicates number of IFNB1 mRNAs. (n = 399 cells in 3 independent experiments) D) Fraction of infected cells expressing >20 IFIT1 mRNAs (black bars) or >10 IFNB1 mRNAs (red bars) at different time periods since the start of phase 3. (n = 399 cells in 3 independent experiments) E) Average viral load of 5xSunTag-EMCV(L^Zn^) infected IFIT1- and IFIT1+ cells at different time periods in infection (relative to the start of phase 3) (n = 243 and 156 IFIT1- and IFIT1+ cells respectively in 3 independent experiments). * indicates p < 0.05 using a two-way ANOVA test. F) Average viral load of 5xSunTag-EMCV(L^WT^) infected, IFIT1- and IFIT1+ cells at different time periods in infection (relative to the start of phase 3) (n = 26 and 106 IFIT1+ and IFIT1-cells respectively in 6 independent experiments). * indicates p < 0.05 using a two-way ANOVA test. G) Number of IFIT1 mRNAs for cells in which infection did or did not progress to phase 3 (n = 118 and 24 phase3+ and phase3-cells respectively in 3 independent experiments) Error bars indicate s.e.m. in all panels

Next, we determined the viral load of cells at different time points in infection, allowing reconstruction of average viral load progression in single cells over time during infection (Fig. 2E). For this, we measured the average EMCV genome FISH signal in infected cells. This analysis revealed that the average vRNA load increased rapidly during the first 6 – 8 h and reached a plateau around 8 h after initial replication, comparable to what is observed when vRNA replication is measured by qPCR (Fig. 2E, S2C). We note that Fig. 2E first shows a significant increase in viral load only at 4h, even though the smFISH-based approach has single-genome detection sensitivity. This detection lag is caused by the very weak EMCV signal in single cell smFISH intensity measurements early in infection combined with minor variations in cellular background fluorescence. Interestingly, cells that activated an antiviral response, as determined by IFIT1 expression, showed a lower average vRNA load than cells that do not express IFIT1 (Fig. 2B,E). Comparison of viral loads between IFIT1+/IFNB1-cells and IFIT1+/IFNB1+ cells was difficult due to the low number of IFIT1+/IFNB1+ cells in the population (Fig. S2D). Importantly, when cells were infected with 5xSunTag-EMCV(L^WT^), instead of 5xSunTag-EMCV(L^Zn^), we observed a similar correlation between viral load and IFIT1 expression, indicating that even in the presence of potent suppression of antiviral signaling, activation of the antiviral response also negatively correlates with viral load (Fig. 2F). We did not detect any IFNB1+ cells upon 5xSunTag-EMCV(L^WT^) infection indicative of the potent antagonism exerted by the L protein and illustrating the benefit of using EMCV(L^Zn^) and IFIT1 as a reporter gene. Notably, the ability to stratify cells using time-lapse microscopy data according to the duration of infection was crucial to reveal the correlation between viral load and innate immune response, as no correlation between viral load and antiviral gene expression levels was observed when infection duration was not taken into account (Fig. S2E), possibly explaining contradictory results on the correlation between viral load and antiviral gene expression in previous studies (Doğanay et al., 2017; Drayman et al., 2019; Fiege et al., 2021; O’Neal et al., 2019; Patil et al., 2015; Zhao et al., 2012).

Previously, using VIRIM on the related picornavirus CVB3, we found that approximately 20% of virus infections arrest before or during replication of the incoming ‘mother’ vRNA (phase 2, see Fig. 1A) (Boersma et al., 2020). For other viruses, abortive infections have been reported to activate the antiviral response more frequently (Drayman et al., 2019). Therefore, we examined IFIT1 expression in cells in which the infection arrested during phase 2. We found that approximately 15% of EMCV infections fail to progress beyond replication of the incoming vRNA. However, abortive infections completely failed to induce IFIT1 expression (Fig. 2G), confirming that viral replication is critical for activation of an innate immune response towards EMCV. A likely explanation for this finding is that insufficient dsRNA for mounting an antiviral response is produced during infections that arrest before or during the first round of replication.

### Real-time visualization of the antiviral response

Results described above show that the average vRNA load in IFIT1+ cells is lower than in IFIT1-cells. These results could indicate that viral replication rates determine the efficiency of antiviral response activation. Alternatively, lower vRNA load in IFIT1+ cells could be the *consequence* of innate immune activation in those cells (i.e. expression of antiviral genes in IFIT1+ infected cells inhibiting viral replication). Determining viral replication rates *before* innate immune activation could help to discriminate between these scenarios; if viral replication rates are already lower in IFIT1+ cells compared to IFIT1-cells before innate immune activation, this would indicate that the differences in vRNA load between IFIT1- and IFIT+ cells are not a *consequence* of innate immune activation, but rather would suggest that slower viral replication *causes* increased innate immune activation. Therefore, we set out to develop a live-cell assay to visualize IFIT1 transcription and virus replication concurrently in single cells. For this, we made use of the PP7 system for fluorescent labeling of single mRNAs (Fig. 3A) (Chao et al., 2008). We inserted an array of 24 PP7 binding sites (PBS) into the endogenous IFIT1 gene along with the SNAP-tag (a protein that allows fluorescent labeling using small molecule dyes) (Keppler et al., 2003) and a puromycin resistance cassette for selection of correctly targeted cells (Fig. S3A). These cells additionally expressed the PP7 coat protein (PCP), which binds with high affinity to the PBS, fused to 3xmCherry and a nuclear localization signal (“PCP-mCherry-NLS”). In this system, transcription of the targeted IFIT1 gene locus results in the synthesis of IFIT1 mRNAs containing the 24xPBS, which are rapidly labeled by PCP-mCherry-NLS, resulting in the appearance of a fluorescent spot at the transcription site (Fig. 3A). As genes are often transcribed by multiple polymerases simultaneously, the transcription site containing nascent RNA is generally brighter than single, ‘mature’ transcripts, allowing discrimination between both types of foci. This real-time IFIT1 transcription imaging system allows live-cell analysis of innate immune activation and enables precise determination of the onset time of IFIT1 transcription, as it measures IFIT1 transcription rather than accumulation of mature IFIT1 transcripts over time, providing an additional advantage over the smFISH-based analysis.

**Fig 3.**
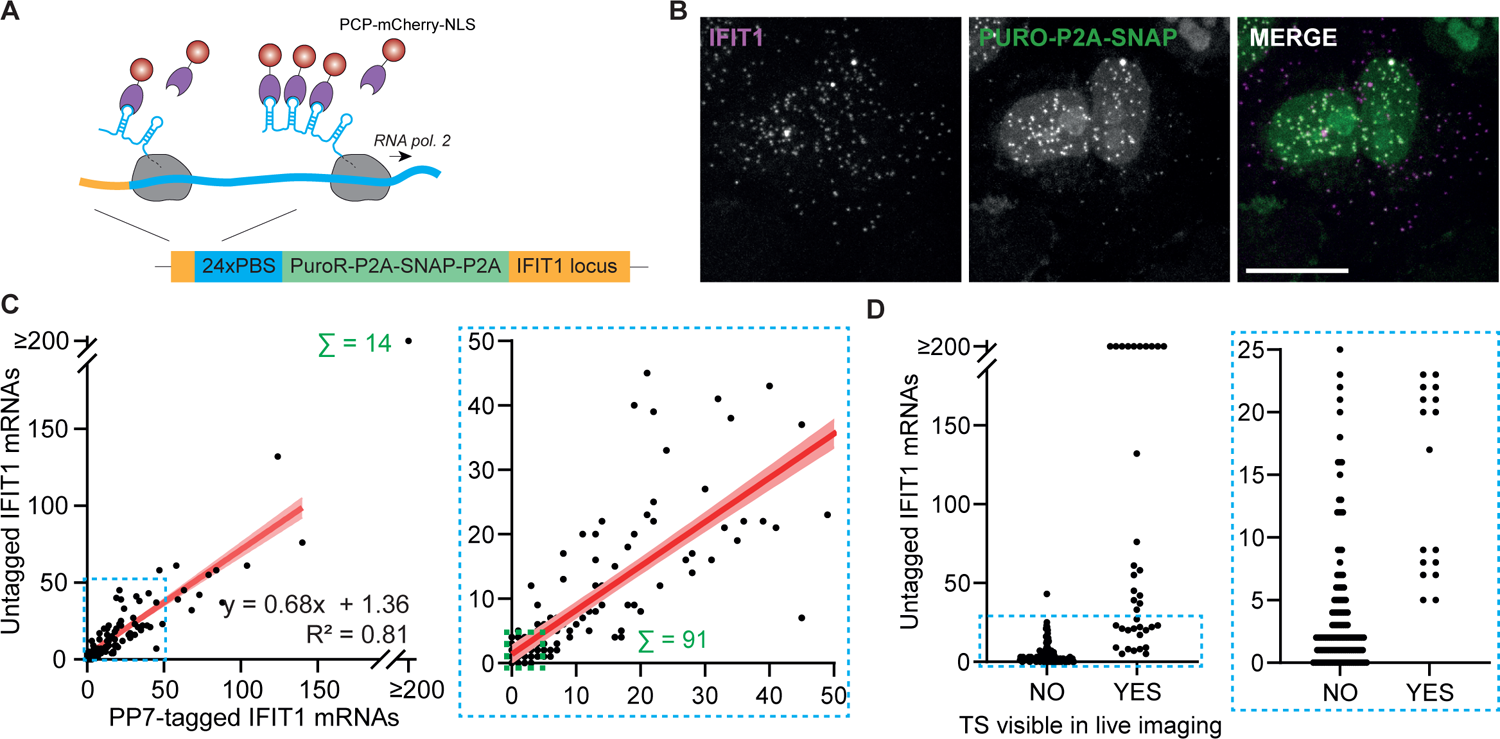
Generation of a live-cell sensor for antiviral response activation. A) Schematic representation of IFIT1 transcription imaging using 24xPBS reporter knockin in the IFIT1 locus. A human RNA polymerase 2 is shown that is transcribing the genetically-modified IFIT1 locus. At the 5’ end, an array of 24 PP7 binding sites is inserted, which, upon transcription, is rapidly labeled by PCP-mCherry-NLS. PBS = PP7 binding site, PCP = PP7 coat protein For panels B), C) and D) 24xPBS-IFIT1 k.i. cells were infected with 5xSunTag-EMCV(L^Zn^) for 16 h and subjected to smFISH using a probe set targeting the IFIT1 coding sequence (which labels mRNAs derived from both the tagged and untagged IFIT1 allele) and a probe set targeting the PURO-P2A-SNAP coding sequence (which only labels mRNAs derived from the tagged IFIT1 allele). B) Representative image of infected 24xPBS-IFIT1 k.i. cell subjected to dual smFISH labeling of both PP7-tagged and untagged IFIT1 mRNAs. Scale bar indicates 20 µm. C) Scatter plot showing the number of PP7-tagged and untagged IFIT1 mRNAs in individual 24xPBS-IFIT1 k.i. cells. Red line is the fit using a linear regression with light red shading indicating the 95% confidence interval of the best fit. Scatter plot on the right represents zoom in of the blue dashed region of the left graph (R^2^ indicates coefficient of determination, n = 177 cells in 3 independent experiments). Multiple data points overlap at the maximum detection level (i.e. 14 cells were detected with over 200 reporter and unlabeled IFIT1 mRNAs, indicated by the green Σ in the left scatter plot) and at base of the graph (i.e. 91 cells had fewer than both 5 reporter and unlabeled IFIT1 mRNAs, indicated with the green box in the right scatter plot) D) Graph shows the number of untagged IFIT1 mRNAs in cells that did or did not develop an IFIT1 transcription site during the 16 h of imaging. Right scatter plot represents zoom in of the blue dashed region in the left plot (n = 38 cells that presented with an IFIT1 transcription site during live imaging and 135 cells did not activate IFIT1 transcription in 3 independent experiments)

To validate that expression of the PP7-tagged IFIT1 allele accurately reports on IFIT1 expression, we performed dual labeling smFISH with one set of probes targeting the tagged IFIT1 allele specifically and a second set (complimentary to the IFIT1 coding sequence) targeting mRNAs originating from both the PP7-tagged and untagged IFIT1 alleles (Fig. 3B). Using this method, we find that the reporter cell line likely has a single integration site of the 24xPBS reporter sequence, because the majority of cells showed only a single transcription site (Fig S3B) (cells in which two 24xPBS reporter transcription sites were detected were likely to be in G_2_ phase of the cell cycle). Moreover, the transcription site labeled by the reporter-sequence specific probes co-localized with IFIT1 specific probes, indicating that the PBS array had integrated into the IFIT1 locus (Fig. 3B). To determine whether the tagged allele was expressed similarly as the untagged allele, we compared the number of mRNAs expressed from both alleles upon viral infection. We find a strong correlation between the expression levels of untagged IFIT1 mRNA and PP7-tagged IFIT1 mRNA (Fig. 3C), demonstrating that tagging of the IFIT1 allele does not alter its expression. Next, we asked whether live-cell analysis of the PP7-tagged IFIT1 allele accurately reported on IFIT1 transcription. For this, we combined live-cell imaging of IFIT1 transcription using the PP7 system with subsequent smFISH of the same cells. We find a strong correlation between cells showing IFIT1 transcription in live-cell imaging and cells showing IFIT1 expression by smFISH (Fig. 3D). Together, these results show that the 24xPBS-IFIT1 locus allows sensitive and accurate live-cell measurements of IFIT1 transcription and can therefore be used as a real-time readout to monitor innate immune activation in single cells.

### Early viral replication rates are lower in cells that activate an antiviral response

While VIRIM allows sensitive, quantitative measurements of viral replication during early infection, late stage infection cannot be readily assessed using VIRIM; the large amount of SunTag protein produced during later stages of infection ultimately sequesters all cellular GFP-STAb, resulting in decreased GFP-STAb labeling of nascent viral proteins and loss of the ability to visualize translating viral genomes. Therefore, we also set out to develop a system that allows simultaneous monitoring of both early and late viral replication. To measure viral replication rates beyond the time of GFP-STAb sequestration, we made use of the previously-developed split-GFP system (Kamiyama et al., 2016). In this system, a small fragment of the GFP protein (GFP beta strand 11; GFP11) is encoded by the virus, while the large remaining fragment of GFP (GFP1-10) is expressed in the cell. Upon infection, GFP11 is synthesized from the viral genome and associates with GFP1-10 to reconstitute fluorescent GFP. While the split-GFP system reports on viral protein levels, the rate of GFP fluorescence increase during infection is dependent on the number of viral genomes that are translated. Therefore, the increase in GFP fluorescence indirectly reports on viral genome abundance and thus on viral replication. Importantly, the split-GFP system lacks the sensitivity of VIRIM during early infection, but reports on viral replication in later stages of infection and is therefore complementary to VIRIM (Fig. 4A,B). To visualize both early and late infection in single cells, we generated GFP11-5xSunTag-EMCV virus and stably expressed the GFP1-10 in the 24xPBS-IFIT1 k.i. cell line expressing GFP-STAb, such that SunTag translation, split-GFP reconstitution and IFIT1 transcription can all be visualized in the same cell (Fig. 4A, Video S2). While both the VIRIM and split-GFP systems have their readouts in the GFP channel, both can be accurately assessed in the same cell, because early in infection the signal originating from the split-GFP is low, allowing readout of the VIRIM signal (i.e. GFP foci), while later in infection (around 3 h after the start of phase 3), the signal originating from the split-GFP system becomes strong enough to detect over the ‘background’ GFP signal originating from GFP-STAb (Fig. 4B,C). After initial detection, split-GFP signal increases for approximately 4 h before reaching a plateau (Fig. 4B,C). Thus, by combining VIRIM with split-GFP imaging, we established a dual-imaging modality that allows both visualization of early viral infection, enabling accurate determination of the start of infection, and measurements of viral replication later in infection.

**Fig 4.**
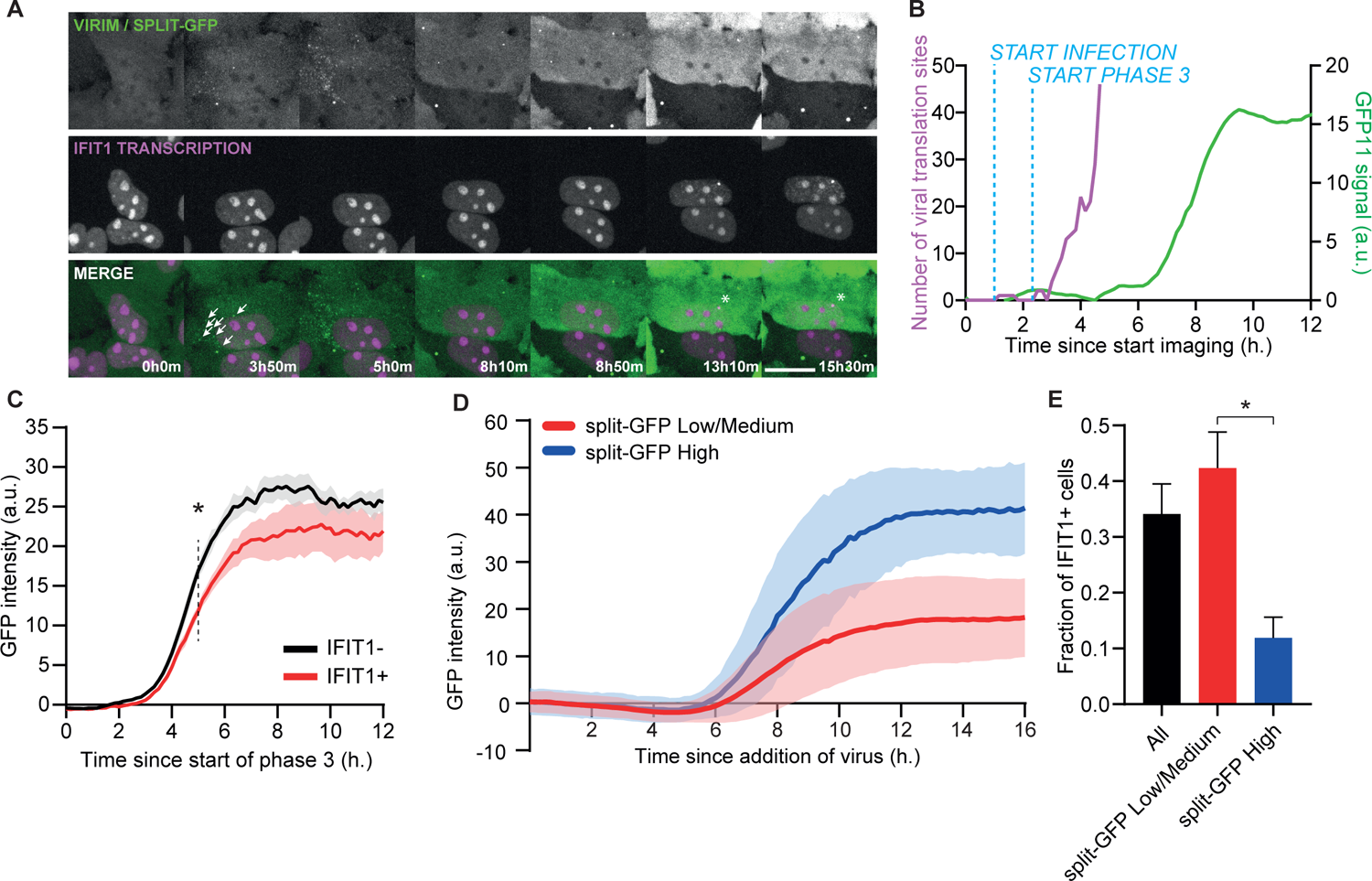
Early viral replication rates are slower in cells that activate an antiviral response. For all panels, 24xPBS-IFIT1 k.i. cells expressing STAb-GFP, GFP(1-10) and PCP-mCherry-NLS were infected with GFP11-5xSunTag-EMCV(L^Zn^) and imaged for 16 h. A) Representative images at different time points from a 16 h movie of cells infected with GFP11-5xSunTag-EMCV(L^Zn^). Top row (green): GFP channel used for VIRIM and at later time points to assess expression levels of the split-GFP system. Middle row (magenta): mCherry channel used for IFIT1 transcription imaging. Bottom row: merge. White arrows at 3h50m time point highlight single viral translation sites that mark the start of phase 3. White asterisk at 13h10 and 15h30 time points indicate IFIT1 transcription site. Scale bar indicates 20 µm. B) Example quantification of GFP11-5xSunTag-EMCV(L^Zn^) replication in a single infected reporter cell based on VIRIM (purple line) and split-GFP signal (green line). By determining the number of viral translation sites in the first hours of infection VIRIM allows determination of the start of infection (appearance of a single viral translation site) and start of phase 3 (first appearance of multiple viral translation sites), which marks the moment of successful replication of the incoming vRNA. Split-GFP signal accumulation can be detected over STAb-GFP signal from approximately 3 h after start of phase 3 onwards. C) Average split-GFP signal accumulation in cells that activate IFIT1 transcription (red line) and cells that do not activate IFIT1 transcription (black line). Line and light shading represent average and s.e.m. of 4 independent experiments (n = 46 IFIT1+ and 117 IFIT1-infections in 4 independent experiments) * indicates p < 0.05 using independent samples T-test at t = 5 h. D) Average GFP intensity time traces of GFP medium/low (red) and GFP high (blue) infections. Line and light shading represent average and s.d. of 3 independent experiments (n: GFP medium/low = 842 infections, GFP high = 129 infections in 3 independent experiments). E) Average fraction of cells that activate IFIT1 transcription in all infections and infections classified as GFP medium/low or GFP high. (n = 3 experiments, error bars = s.e.m.) * indicates p < 0.05 using paired samples T-test.

Next, we compared split-GFP intensity time traces of cells that activate IFIT1 transcription with cells that do not initiate IFIT1 transcription. This analysis revealed that split-GFP signal increased slower in IFIT1+ cells (Fig. 4C,S4A,B), indicative of slower replication rates. This difference in split-GFP accumulation rate is already apparent at 5 h after start of phase 3 (Fig. 4C), well before IFIT1 transcription typically initiates (See fig.2D). These findings indicate that the lower average viral load in IFIT1+ cells compared to IFIT1-cells is not caused by virus-induced antiviral gene expression limiting viral replication, but rather indicates that the rate of viral replication itself affects the efficiency of antiviral response activation.

Although the average viral replication rate, as determined from GFP intensity time traces, correlated with the ability of cells to activate an antiviral response, the predictive power at the single cell level was modest. Therefore, we performed more in-depth analysis of split-GFP intensity time traces to better predict antiviral response activation for individual cells based on viral replication rates (i.e. on split-GFP expression dynamics). We developed an automated analysis pipeline to measure split-GFP intensities and performed unbiased clustering on the resulting intensity time traces. Using this approach, we were able to identify a group of infections (∼15% of all infections), characterized by rapid split-GFP signal accumulation and high split-GFP plateau intensities (Fig. 4D, S4C). In this group of infections with high split-GFP signal, the vast majority of cells did not activate the antiviral response (Fig. 4E), demonstrating that in cells exhibiting high viral replication rates antiviral response activation is highly inefficient. Together, these findings indicate that heterogeneity in viral replication rates shape the host cell’s ability to activate the antiviral response and induce antiviral gene expression.

### Efficient activation of the antiviral response occurs during a defined time window

The considerable time lag between the first round of viral replication (i.e. replication of the incoming vRNA) and the initiation of IFIT1 transcription (See fig. 2D) suggests that activation of the antiviral response does not occur efficiently early in infection. Therefore, we wished to more precisely assess when innate immune activation occurs throughout infection. We determined the onset of IFIT1 transcription and found that over 90% of the IFIT1+ cells activate IFIT1 transcription between 5 and 10 h after the first viral dsRNA was present in the cell (Fig. 5A). These results not only confirm that innate immune activation does not occur early in infection, but also reveal that activation of IFIT1 transcription in the final hours of infection (>10 h) is rare. Next, we assessed how the moment of IFIT1 transcription activation relates to the strength of IFIT1 transcription (as determined by the IFIT1 transcription site intensity). We found that IFIT1 transcriptional activity was on average ∼3-fold higher when IFIT1 transcription was activated at 5 h vs 10 h after initiation of vRNA replication (Fig. 5B), indicating that the antiviral response is most efficient when activated early in infection. Notably, a CMV-driven reporter gene showed constant transcription rates throughout infection (Fig. S5), indicating that reduced IFIT1 transcriptional activation at later time points in infection was not due to global virus-induced transcriptional inhibition in these cells. Together these findings demonstrate that the probability of antiviral response activation varies during infection and is most efficient during a defined ‘window of opportunity’ around 5 h after initial replication. An inability to activate the dsRNA sensing pathway during this window of opportunity may cause a blunted or absent antiviral response.

**Fig. 5.**
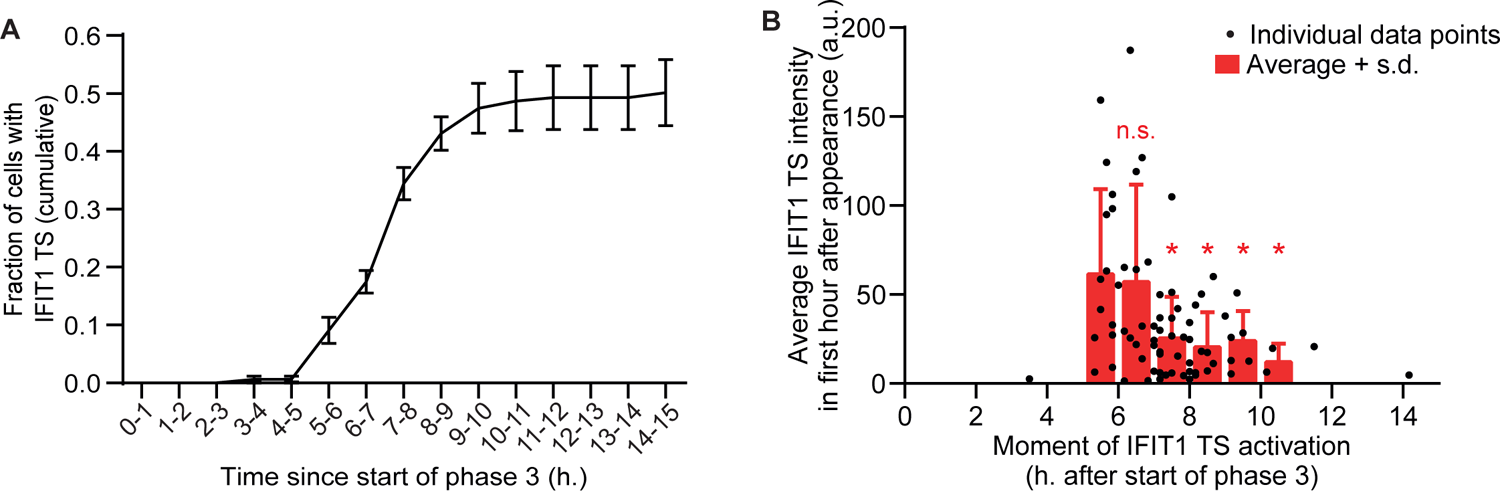
Activation of the antiviral response occurs most efficiently during a brief time window in infection. For both panels, 24xPBS-IFIT1 k.i. cells stably expressing STAb-GFP, PCP-mCherry-NLS were infected with 5xSunTag-EMCV(L^Zn^) and imaged for 16 h. A) Cumulative fraction of cells that have activated IFIT1 transcription at different time points since the start of phase 3. Line indicates average of 4 experiments, error bars = s.e.m. (n = 158 infections). B) Scatter plot showing the moment of IFIT1 transcription activation and average IFIT1 transcription site intensity in the first hour. Red bars indicate average IFIT1 transcription site intensity in different time bins, error bars indicate s.d. (n = 76 cells in 4 independent experiments) Red * indicates p < 0.05 using independent samples T-test compared to 5 – 6 h time period. Red n.s. indicates non significant.

### Differential IRF3 nuclear translocation explains heterogeneous antiviral responses

The absence of antiviral gene expression in a subset of cells may result from inefficient activation of the dsRNA sensing pathway. To investigate this possibility, we set out to monitor the dsRNA sensing pathway through visualization of nuclear translocation of the IRF3 transcription factor, a key downstream event in the activation of the dsRNA sensing pathway. We tagged the endogenous IRF3 protein with BFP using the CRISPR/Cas9 system (Fig. S6A). The resulting IRF3-BFP cell line showed similar innate immune activation as control cells (Fig. S6B), indicating that the efficiency of antiviral response activation is not affected by BFP introduction in the IRF3 locus. Imaging fluorescent IRF3-BFP translocation in the 24xPBS-IFIT1 k.i. cell line (Fig. 6A, Video S3), revealed that 1) efficient IRF3 nuclear translocation is almost exclusively observed in cells that activate IFIT1 transcription (Fig. 6B,C) and 2) IFIT1 transcription is typically activated very shortly after IRF3-BFP nuclear translocation (on average 15 min after translocation) (Fig. 6D). These findings indicate that the lack of IFIT1 expression in a subset of cells is caused by inefficient IRF3 nuclear translocation, likely as a consequence of poor activation of the dsRNA sensing pathway, rather than of cell-to-cell variation in the ability of IRF3 to transcriptionally activate the IFIT1 gene (for instance because of differences in the epigenetic status of the IFIT1 locus). Moreover, since IRF3 nuclear translocation temporally coincides with IFIT1 transcriptional activation (Fig. 6B,D), these results show that the lag between initial viral replication and IFIT1 expression (see Fig. 2D, 5A) is caused by inefficient activation of the viral sensing pathway, rather than by slow transcription activation of IRF3 target genes.

**Fig. 6.**
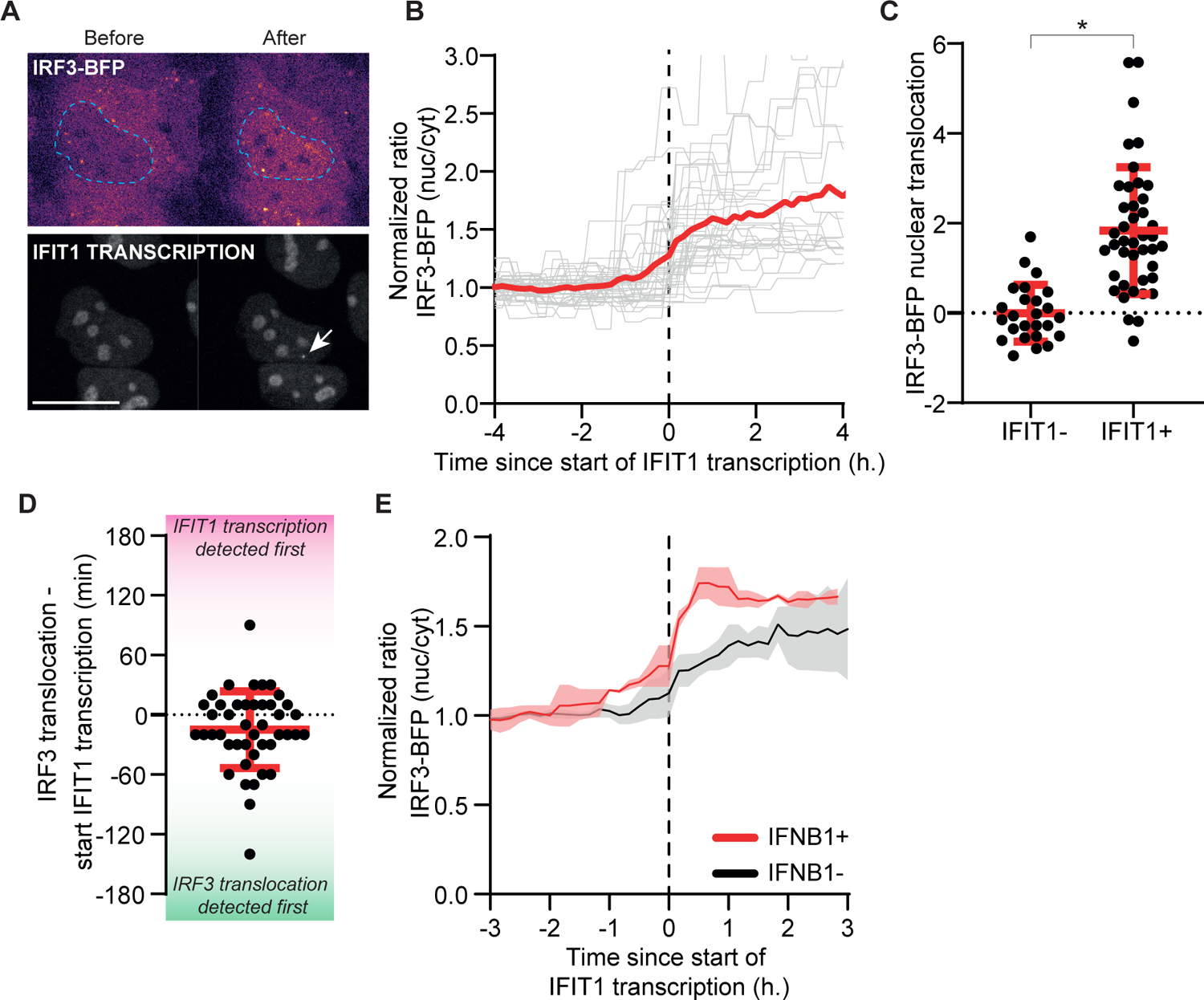
Dynamics of IRF3-BFP nuclear translocation control heterogeneous outputs of the antiviral response. For all panels, IRF3-BFP and 24xPBS-IFIT1 k.i. cells stably expressing STAb-GFP and PCP-mCherry-NLS were infected with 5xSunTag-EMCV(L^Zn^) and imaged for 16 h. In panel E, live-cell imaging was followed by fixation and smFISH with probes targeting IFNB1 mRNA and imaged cells were retrieved to link IRF3-BFP nuclear translocation efficiency to IFNB1 expression status. A) Representative image of IRF3-BFP localization (top row) before and after IFIT1 transcription activation (bottom row). Note that IRF3-BFP has translocated to the nucleus upon IFIT1 transcription activation. IRF3-BFP signal is pseudo-colored to aid visualization. Blue dotted line in top row indicates outline of the nucleus. White arrow in bottom row indicates IFIT1 transcription site. Large nuclear foci represent PCP-mCherry-NLS accumulation in nucleoli. Scale bar indicates 20 µm. B) Normalized nucleocytoplasmic ratios of IRF3-BFP over time. Time traces of single cells were aligned to the onset of IFIT1 transcription (set as t = 0). See method section for explanation about the quantification of normalized nucleocytoplasmic ratio. Red line indicates average of all traces and grey lines represent individual traces (n = 21 cells in 2 experiments) C) Quantification of IRF3-BFP nuclear translocation efficiency in IFIT1- and IFIT1+ cells. See method section for explanation about the quantification of translocation efficiency. Dots = measurements in individual cells, red lines and error bars indicate average ± s.d. (IFIT1- = 25 cells, IFIT1+ = 42 cells in 3 independent experiments) * indicates p < 0.05 using independent samples T-test. D) Time between moment of IFIT1 transcription activation and IRF3-BFP nuclear translocation. Positive values (pink shaded area) indicates start of IFIT1 transcription is detected before IRF3 nuclear translocation, whereas negative values (green shaded area) indicates IRF3 nuclear translocation was observed before IFIT1 transcription activation. Red lines and error bars indicate average ± s.d. (n = 44 cells in 3 independent experiments) E) Normalized nucleocytoplasmic ratios of IRF3-BFP in IFNB1+ (red line) and IFNB1-(black line) cells aligned to the onset of IFIT1 transcription. Solid line represents average from 3 experiments; shaded area indicates s.e.m. (total n = 16 and 29 cells for IFNB1+ and IFNB1-respectively in 3 independent experiments)

Although both IFIT1 and IFNB1 transcription can be triggered upon detection of viral dsRNA, IFNB1 expression is limited to a subset of IFIT1+ cells, indicating that viral sensing can result in diverse antiviral responses. However, the underlying causes of such qualitatively distinct antiviral responses are largely unknown. Intriguingly, when examining IFNB1 levels by smFISH after live-cell analysis of IRF3-BFP nuclear translocation, we found that IFNB1+ cells showed a more pronounced IRF3-BFP nuclear translocation than IFNB1-cells (Fig. 6E). This finding indicates that signaling strength of the dsRNA sensing pathway (reflected by the extent of IRF3 nuclear translocation) differs among infected cells, and suggests that IFNB1 expression requires stronger pathway activation compared to IFIT1 expression. This conclusion is consistent with our finding that IFNB1 expression is mostly observed in cells with very high IFIT1 expression (Fig. 1F). Together, these results show that the signaling that leads to IRF3 nuclear translocation is the rate-limiting step in antiviral gene expression and that differential IRF3 activation in response to infection likely explains the heterogeneity in transcriptional response to viral infection.

## Discussion

Detection of intracellular viral infection and initiation of antiviral gene expression are key steps required for successful elimination of viral infection. By inhibiting antiviral processes of the host cell, viruses engage in a competition with the host cell, of which the outcome varies, resulting in sporadic activation of antiviral gene expression in infected cells. However, the parameters that determine whether an infected cell is able to launch a successful antiviral response are poorly understood. Using a combination of VIRIM with sensitive real-time readouts of the endogenous antiviral response, we were able to generate a detailed, quantitative characterization of the parameters that affect antiviral response activation, which provided several molecular insights into the origin of heterogeneity in antiviral response activation. We show that viral replication rates during early infection determine the efficiency of antiviral gene expression and thereby contribute to heterogeneity in the antiviral response (Fig. 2,4). Moreover, we find that the probability of activating the antiviral response varies throughout infection, with most efficient activation during a defined time window ∼5-7 h after the initial viral replication (Fig. 5A,B). The moment of antiviral response activation during infection was found to correlate with the strength of activation (Fig. 5B), indicating that the timing of dsRNA sensing is an important factor contributing to heterogeneity in antiviral gene expression. Finally, by measuring IRF3 nuclear translocation, we show that IRF3 translocation (i.e. dsRNA sensing pathway activation) is the rate-limiting step in antiviral gene expression, and that quantitative differences in signaling strength of the dsRNA sensing pathway are associated with a qualitatively distinct pattern of antiviral gene expression (Fig. 6). Together, these results reveal that quantitative aspects of viral replication and host sensing shape the outcome of the antiviral response at multiple levels.

Previous studies have examined whether variations in viral load affect antiviral gene expression, but, in contrast to our work, often failed to detect such a correlation (Doğanay et al., 2017; Drayman et al., 2019; O’Neal et al., 2019; Patil et al., 2015). These studies frequently involved fixed-cell, single time point measurements to assess viral load and antiviral gene expression. Such measurements are, however, unable to take into account the variation in the onset time of viral infection and replication. Here, using the VIRIM technology, we were able to image early stages of infection, which revealed considerable cell-to-cell variation in the moment of infection and in the time between infection and completion of initial replication. These variable parameters of infection result in considerable variation in the amount of time that dsRNA has been present in each infected cell at the moment of cell fixation (Fig. S2A). Correcting for this variation, our data demonstrate that an antiviral response is preferentially activated in cells with a lower viral load (Fig. 2E, 4C), an effect that could not be detected without temporal information on early infection (Fig. S2E), and providing a plausible explanation why previous studies failed to detect a similar relationship between viral load and antiviral gene expression.

Why does the viral replication rate inversely correlate with antiviral response activation? Higher replication rates result in a higher concentration of dsRNA in the cell, providing more viral ligands that can trigger immune activation. Counterintuitively, we find that faster replication is associated with less efficient antiviral response activation (Fig. 4C). A likely explanation for this paradox is that in cells in which infection proceeds faster, viral proteins accumulate more rapidly and can impair host antiviral response pathways more effectively early in infection. Rapid inhibition of these host pathways limits the ability of the host to mount an antiviral response once sufficient viral dsRNA is formed for efficient dsRNA sensing, outweighing the increased probability of dsRNA detection by the host. Although the prime host antagonist of EMCV, i.e. the Leader (L) protein, is inactivated in the recombinant EMCV(L^Zn^) virus that was used in most of our experiments, additional EMCV proteins have been implicated in suppressing the dsRNA sensing pathway (Han et al., 2021; Hato et al., 2007; Huang et al., 2017; Li et al., 2019). Importantly, a lower viral load was also observed in IFIT1+ cells upon infection with EMCV(L^WT^) (Fig. 2F), indicating that heterogeneity in viral replication similarly affects the efficiency of antiviral response activation when the dsRNA sensing pathway is more efficiently inhibited.

Little is known about the factors that determine viral replication rate and how such factors could lead to variation in replication rates among infected cells. Both host-cell intrinsic and virus-intrinsic factors may affect viral replication rates (Guo et al., 2017; Jones et al., 2021). Previous work using a fluorescent-reporter expressing poliovirus found that variation in viral replication rates, as determined by the maximum rate of fluorescence increase, was of viral-intrinsic origin, and does not result from differences between host cells (Guo et al., 2017). If these findings hold true for other viruses, then virus-intrinsic variability in replication rates are likely relevant for shaping the antiviral response. The molecular basis for such virus-intrinsic differences in replication rates would be an important future topic of research.

We demonstrate that antiviral gene transcription is not efficiently activated until ∼5 h after the first round of vRNA replication, (Fig. 2D, 5A). Since we show that nuclear translocation of IRF3 (and thus activation of the dsRNA sensing pathway) temporally correlates with antiviral gene transcription, these results indicate that the dsRNA sensing pathway is not activated during the first hours of infection. The inability to activate this pathway early in infection may be caused by the relatively low concentration of dsRNA present in the cell at this stage, which would suggest that dsRNA sensing mechanisms are relatively insensitive and require a large number of dsRNA molecules to become activated. Such low sensitively could be caused by inefficient detection of dsRNA molecules by dsRNA sensors (e.g. RLRs), or may result from inefficient relay of dsRNA detection signals to downstream activation of the innate immune pathway (i.e. IRF3 translocation). Alternatively, masking of viral dsRNA, for example through formation of replication organelles, could prevent dsRNA detection during the first few hours of infection, until the amount of dsRNA exceeds the shielding capacity of these organelles (Albulescu et al., 2015; Belov and van Kuppeveld, 2012). Irrespectively, our findings show that once dsRNA sensing induces IRF3 nuclear translocation, antiviral gene expression occurs reliably and fast (Fig. 6B,C,D), suggesting that the early steps in innate immune pathway activation represent the bottleneck for activation. Although a relatively low sensitivity of the dsRNA sensing pathway early in infection provides a head start for viruses, it may have evolved to prevent spurious immune activation by endogenous dsRNA ligands in uninfected cells, protecting cells and tissues from an inappropriate inflammatory response. In this context, it is interesting to note that differences in IRF3 nuclear translocation efficiency resulted in distinct transcriptional responses, with IFNB1 expression being associated with higher pathway activation (Fig. 6E). These findings suggest that signaling strength of the dsRNA sensing pathway dictates which set of antiviral genes are expressed upon detection of viral presence. Since IFN is a potent activator of inflammation, the higher dsRNA sensing pathway activation required for IFN production may further help to limit tissue damage due to inappropriate innate immune activation upon sensing of endogenous dsRNA. Moreover, the ability to modulate the set of antiviral genes that are transcribed dependent on the activity of the dsRNA sensing pathway may allow graded or functionally distinct responses to various types of infection.

We also find that IFIT1 transcriptional activity is reduced when innate immune activation occurs late in infection (between 8-10 h) (Fig. 5B). As the transcriptional capacity of EMCV(L^Zn^) infected cells does not globally decrease (Fig. S5), this finding suggests that when activated at these later time points the antiviral response is already partially compromised, likely by viral antagonism. Together, inefficient antiviral pathway activation during both early and late infection results in a brief time period during which the antiviral response can be efficiently activated. The finding that most efficient activation of the antiviral response occurs during a limited time window indicates that the timing of dsRNA sensing is an important factor that contributes to quantitative differences in antiviral gene expression among infected cells.

In summary, by using new techniques that allow real-time imaging of both virus infection and antiviral response activation we show that the dynamics of viral replication and host cell sensing shape heterogeneity in the antiviral response.

## Acknowledgements

We thank Tim Hoek for help with the logistic growth model simulations. We thank members of the Tanenbaum lab and van Kuppeveld lab for helpful discussions. This work was financially supported by an ERC starting grant to M.E.T. (EU/ERC-677936 RNAREG), a NWO klein-2 grant (OCENW.KLEIN.344) to M.E.T. and F.J.M.v.K, the Howard Hughes Medical Institute through an international research scholar grant to M.E.T. (HHMI/IRS 55008747), and a NWO VICI grant to F.J.M.v.K. (91812628). We would also like to thank the Friends of the Hubrecht Institute for financial support. L.J.M.B., M.M., H.R., S.B. and M.E.T. were supported by the Oncode Institute, which is partly funded by the Dutch Cancer Society (KWF).

## Author contributions

Conceptualization, L.J.M.B., F.J.M.v.K., and M.E.T.; Methodology, L.J.M.B., M.M., J.G.S., F.J.M.v.K., and M.E.T.; Validation, S.B.; Formal Analysis, L.J.M.B., M.M., J.G.S.; Investigation, L.J.M.B., J.G.S., and H.H.R.; Data Curation, L.J.M.B., M.M., J.G.S. and H.H.R.; Writing – Original Draft, L.J.M.B., F.J.M.v.K., and M.E.T.; Writing – Review & Editing, all authors; Visualization, L.J.M.B; Supervision, F.J.M.v.K. and M.E.T.; Funding Acquisition, F.J.M.v.K. and M.E.T

## Methods

### Cell lines

HeLa, HEK293T and BHK-T7 cells were cultured in DMEM (GIBCO) supplemented with 10% FCS (Sigma-Aldrich) and 1% PenStrep (GIBCO). HeLa MDA5 and MAVS k.o. cell lines were previously established (Melia et al., 2017; Schuster et al., 2017). Cells were cultured at 37°C and 5% CO_2_. Cell lines used in this study were routinely tested for presence of mycoplasma.

### Chemicals

The following inhibitors were used in this study: Dipyridamole (DiP, 25μM, Sigma-Aldrich), MRT67307 (MRT, 1μM, Sigma-Aldrich), Tofacitinib (TOFA, 1μM, Sigma-Aldrich). All inhibitors were added to cells 30 min prior to virus addition.

### Virus design and production

5xSunTag-EMCV was produced as described previously (Boersma, et al., 2020). Briefly, a 5xSunTag array was introduced in the infectious pM16.1 cDNA clone of the Mengovirus strain of EMCV (kindly provided by A. Palmenberg) (Duke and Palmenberg, 1989). The array was introduced after codon 6 of the Leader protein and was followed by a 3C(D) cleavage sequence (VFETQG) to allow release from the viral polyprotein.

For GFP11-5xSunTag EMCV, the GFP11 coding sequence was inserted upstream of 5xSunTag array using Gibson assembly with an annealed oligo pair. No additional cleavage sequence was introduced between the GFP11 and 5xSunTag sequence. To enable infectious RNA production in T7 expressing BHK cells, a T7 terminator sequence was introduced downstream of the viral polyA sequence using Gibson assembly with annealed oligos.

Virus stocks were generated by either transfecting purified in vitro transcribed viral RNA (HiScribe, NEB) or transfecting the infectious cDNA clone containing plasmid in BHK-T7 cells. The day after transfection medium was refreshed and 2-4 days after transfection, when a significant cytopathic effect was observed, remaining cells and the supernatant were harvested and subjected to 3 cycles of freeze-thawing. Cellular debris was cleared by centrifugation and supernatants were collected. Virus titers were determined by end-point titration and viral RNA was extracted from particles to verify the insert sequence by RT-PCR and sanger sequencing.

### Cell culture, cell stimulation and infection for live imaging

One day prior to imaging cells were seeded on a 96-well glass bottom plate (Matriplates, Brooks life science systems) such that cells were at ∼80% confluency at the start of imaging. 30 min Before start of imaging, medium was replaced by Leibovitz’s L15 medium (GIBCO) supplemented with 10% FCS and 1% PenStrep. An MOI of ∼1 was used for all imaging experiments.

### Reporter cell line generation

#### (Lentiviral) transduction

GFP-STAb, PCP-mCherry-NLS, GFP(1-10)-P2A-PuroR were introduced into cells using lentiviral infection. For this, pHR-based lentiviral plasmids containing these transgenes were transfected into HEK293T cells together with pMD2G and psPAX2 helper plasmids using Fugene (Promega). After 2 days, viral supernatant was passaged over a 0.45μm filter to remove cellular debris and polybrene (2μg/ml, Santa Cruz) was added before transferring the virus-containing supernatant to recipient HeLa cells. Two days after virus transfer, medium was replaced and after two passages single cells from the polyclonal cell population were sorted in 96-well plates by FACS. Cells with GFP-STAb expression were selected to have a similar GFP intensity as a previously established U2OS-GFP-STAb monoclonal cell line that is routinely used for translation imaging in our lab (Yan et al., 2016). Cells with PCP-NLS-mCherry expression were sorted for low mCherry fluorescence. GFP(1-10)-P2A-PuroR transduced cells were treated with puromycin (1μg/ml, ThermoFischer) 3 days before sorting. After expansion, correct monoclonal cell lines were selected based on expression levels of the transgenes.

To generate a doxycycline-inducible, CMV-promotor driven, 24xPBS transcription reporter, PCP-mCherry-NLS expressing HeLa cells were infected with lentiviral particles to express TetR-2A-HygroR. Two days after infection cells were selected for hygromycin resistance using 200μg/ml hygromycin (Invivogen). A pcDNA3-based plasmid containing the CMV-TetOn-24xPBS transcription reporter was transfected into the TetR expressing HeLa cells using Fugene according to the manufacturers’ instructions. Two days after transfection, medium was replaced and cells with stable integration of the plasmid were selected using zeocin for 2 weeks (0.4mg/ml, Invitrogen). Surviving cells were then sorted as single cells in 96-well plates and expanded to generate monoclonal cell lines. Individual clones were screened using doxycycline stimulation (1μg/ml, Sigma-Aldrich): appropriate clones were selected that had no PP7 transcription site prior to doxycycline addition and which presented a transcription site in the nucleus after doxycycline addition.

### CRISPR-Cas9 genome editing

CRISPR-Cas9 genome editing was used to introduce the 24xPBS transcription reporter and BFP sequence into the IFIT1 and IRF3 locus, respectively. The dsDNA donor template for homology-dependent repair was created by excision of the donor sequence from a plasmid using the Sap1 restriction enzyme. The ends of the donor sequence contained 300bp homology to the genomic sequence of the IFIT1 and IRF3 locus. The 24 PBS hairpins in the reporter were re-designed to remove stop codons from the coding sequence and a P2A-PuroR-P2A-SNAP-tag-P2A cassette was added to the reporter in frame with the downstream IFIT1 coding sequence for selection purposes.

Guide RNA (gRNA) sequences were ligated in a Cas9 plasmid (PX459), that was linearized with Bbs1, and from which the puromycin resistance gene was removed. The following guide sequences were used: IFIT1 5’-TGATTTAGAAAACAGAGTC-3’, IRF3 5’-CATGGATTTCCAGGGCCCTG-3’. The Cas9-guide construct was transfected together with linearized donor template and two days after transfection medium was replaced. The plasmid encoding Cas9 and the gRNA construct were transfected together with the linearized donor template. Two days after transfection medium was replaced.

For the 24xPBS IFIT1 k.i., cells were then stimulated with 100U/ml recombinant IFNα2 (Sigma-Aldrich) to induce transcription from the IFIT1 loci, resulting in expression of the puromycin resistance gene which was integrated into the IFIT1 locus along with the PBS array. After 24h, cells were continuously cultured in the presence of puromycin while receiving fresh IFN-containing medium once every two days to maintain expression of the puromycin resistance gene. Surviving cells were expanded and sorted as single cells in 96-well plates to generate monoclonal cell lines. Genomic DNA of expanded clones was extracted using proteinase K digestion and correct integration of the 24xPBS reporter was confirmed by PCR amplification and sequencing of the edited allele. For the IRF3-BFP k.i. cells no selection was performed prior to single cell sorting by FACS. Monoclones were expanded and screened for BFP expression under the microscope. Correct integration of the BFP sequence was confirmed by PCR amplification of the edited locus and sequencing.

### Single molecule fluorescence in situ hybridization (smFISH)

smFISH was performed according to protocols described previously (Gaspar, et al., 2018; Lyubimova et al., 2013).

### smFISH probe generation

Stellaris probe designer (https://www.biosearchtech.com/support/tools/design-software/stellaris-probe-designer) was used to design probes targeting IFIT1, IFNB1 and Puro-P2A-SNAP (part of the 24xPBS transcription reporter) mRNA or EMCV vRNA. The probe sets for IFIT1, IFNB1, Puro-P2A-SNAP and EMCV contained 48, 31, 36 and 48 unique target sequences respectively. 20-Mer oligonucleotides were ordered from integrated DNA technologies (IDT) and pooled (sequences are listed in table 1). All oligonucleotide probes targeting a single RNA were combined and labeled with ddUTP-coupled Atto-488, Atto-565 or Atto-633 dyes (AttoTec) using Terminal deoxynucleotidyl Transferase (TdT) as described previously (Gaspar et al., 2018). Fluorescent probes were purified by EtOH precipitation, washed to remove unlabeled probes and resuspended in nuclease-free water. Concentration of labeled probes and labeling efficiency was determined by UV/Vis spectroscopy.

**Table 1:**
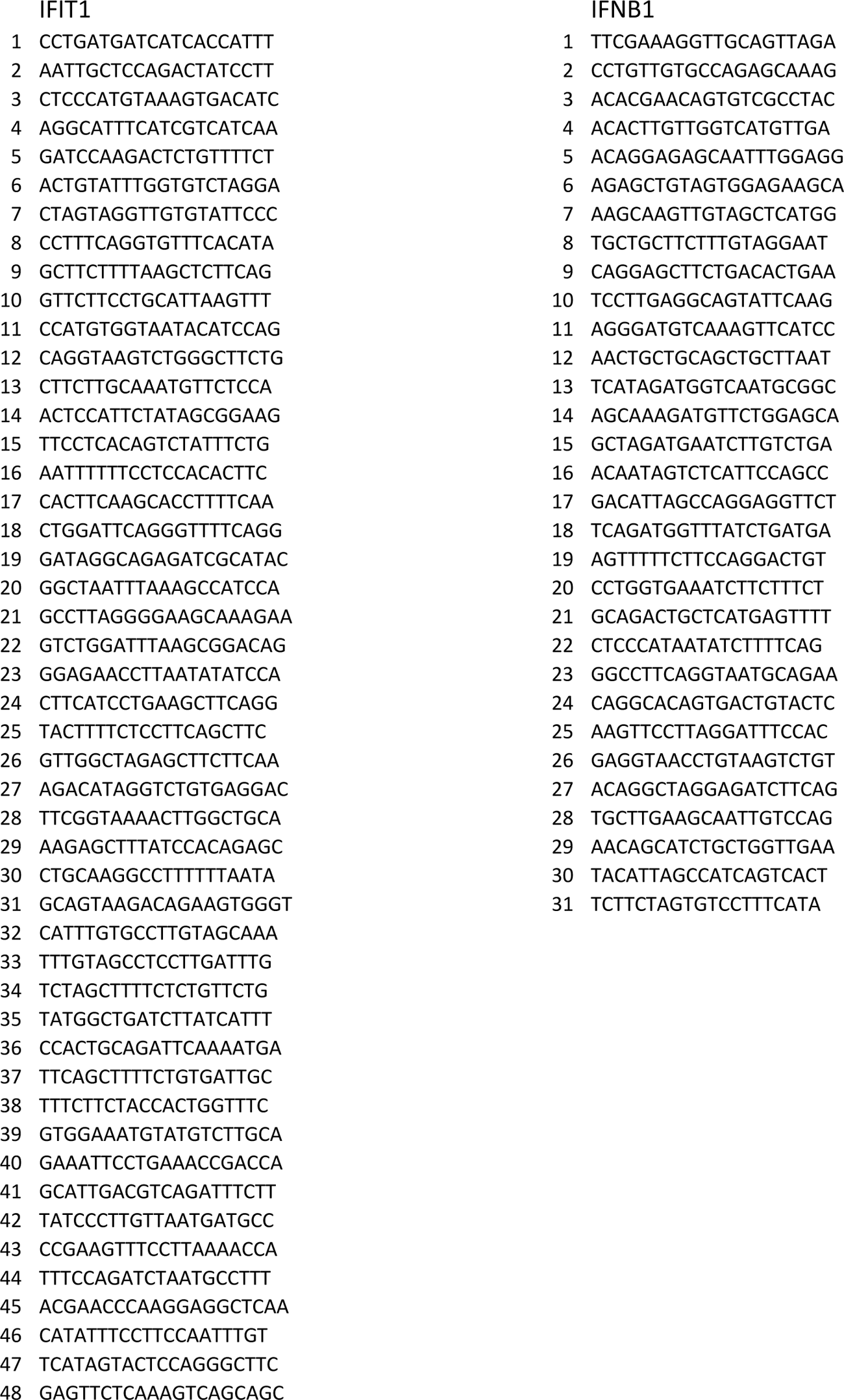
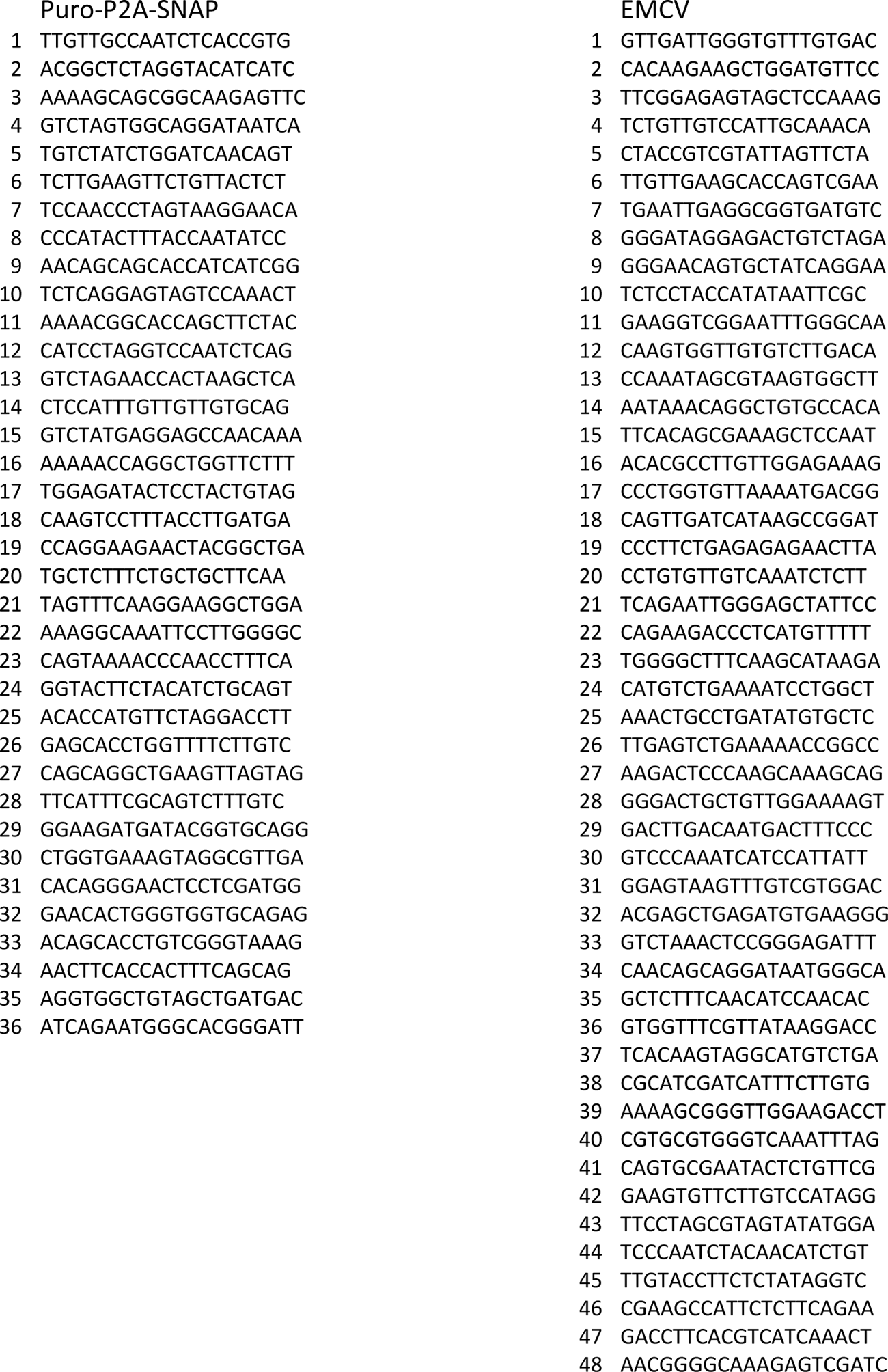
Sequence of smFISH probe sets

### smFISH staining procedure

Cells on 96-well glass bottom plates were washed in PBS0 and fixed in 4% formaldehyde (Electron Microscopy Sciences) for 5 min at RT. After fixation, cells were washed 3x in PBSO and permeabilized in 100% EtOH for 30 min on ice followed by two 15 min washes in wash buffer (2xSSC, 10% formamide in DEPC-treated water) at room temperature (RT). Labeled smFISH probes were diluted to 10nM in hybridization buffer (1% dextran sulfate, 2xSSC, 10% formamide in DEPC-treated water) and hybridization was performed in a sealed, dark container at 37°C for 16 h. Unbound smFISH probes were removed by two 1 h washes in wash buffer at 37°C and a 15 min wash at RT. Samples were stored and imaged in imaging buffer (10mM Tris pH8, 2xSCC, 0.4% glucose, supplemented with glucose oxidase (Sigma-Aldrich) and catalase (Sigma-Aldrich)). Imaging was performed within 3 days after probe hybridization.

### Growth curve and quantitative RT-PCR (qPCR)

HeLa cells expressing STAb-GFP were seeded at 10k cells per well in flat-bottom 96-well plates. Cells were infected on the next day for 30 min with 5xST-EMCV(L^Zn^) at a multiplicity of infection of 5. Medium was refreshed and at indicated time points post infection and plates were either freeze-thawed 3 times to lyse the cells for titration or lysed using RNA lysis buffer for isolation of total RNA for qPCR. Supernatants were titrated with endpoint titration assays to determine viral titers. qPCR was performed to determine the amount of viral RNA copies and expression of IFIT1 and IFNB1 at different time points post infection. For this total RNA was isolated using Nucleospin RNA kits (Machery-Nagel). Subsequently, reverse transcription was performed on the isolated RNA using random hexamer primers and TaqMan reverse transcriptase (Thermo-Fisher Scientific). cDNA was subjected to qPCR with specific primers for IFIT1, IFNB1, EMCV vRNA and Actin. Relative levels of IFIT1 and IFNB1 mRNA and EMCV vRNA were normalized to Actin expression. Results are derived from three experiments and per experiment every time point was determined from the average of three technical replicates.

### Microscopy

All fluorescence microscopy was performed on a Nikon TI2 inverted microscope equipped with a Yokagawa CSU-X1 spinning disk and a Prime 95B sCMOS camera (Photometrics). Imaging was performed using a 60x 1.40 NA oil objective. Image acquisition was performed using NIS Elements software and making use of the ‘perfect focus system’ to correct for Z drift during time lapse imaging experiments. The microscope was equipped with a temperature-controlled incubator and imaging was performed at 37°C for live-cell experiments or at RT for smFISH samples.

### Microscopy acquisition settings

For smFISH samples that were not previously subjected to live-cell imaging, a random position in the center of the well was selected and a large field of view was constructed by imaging 4×4 neighboring imaging fields. Approximately 15 Z-slices at 0.5 μm interval that covered the entire cell were acquired for IFIT1, IFNB1 and Puro-P2A-SNAP smFISH labeled cells and a single Z slice at the center of the cell was acquired for EMCV FISH labeling. IFNB1 and IFIT1 smFISH were acquired using a 50 ms exposure times; EMCV was acquired with an exposure time of 70 ms.

For live-cell imaging experiments that were not followed by smFISH, random non-overlapping positions were selected. Movies with live-cell reporters were acquired with a 5 min time interval between frames with the exception of the BFP channel in experiments involving the IRF3-BFP k.i. cell line. Because of the high laser power required to obtain sufficient BFP signal, we limited the imaging interval to 1 frame per 30 or 60 min to reduce phototoxicity (Initial experiments were performed at 1 frame per 60 min to minimize phototoxicity, in later experiments 1 frame per 30 min was found to be equally well tolerated by cells). Typically, 10 Z-slices at 0.8μm interval were acquired for GFP and mCherry channels, whereas a single Z-slice was acquired for the BFP channel in experiments involving the IRF3-BFP k.i. cell line. Signal in the red channel (561 nm laser), used for imaging of IFIT1 transcription, was acquired using a 50 ms exposure time; signal in the green channel (488 nm laser), used for VIRIM and split-GFP imaging, was acquired with an exposure time of 70 ms; signal in the blue channel (405 nm laser), used to visualize IRF3-BFP localization, was acquired using a 50 ms exposure time.

For live-cell imaging experiments that were followed by smFISH, positions were selected for live imaging in a pattern that could be retrieved. For this, a series of consecutive field of views was selected starting from the edge of the imaging well. Images were taken at a 10 min interval. After completion of the live imaging experiment, the imaging plate was gently removed from the microscope stage and cells were immediately washed and fixed. After completion of smFISH staining protocol, positions were retrieved by navigating to the first field of view at the edge of the imaging well and imaged again to visualize smFISH labeling.

### Post-acquisition data processing

Maximal intensity projection for all Z-slices were generated using NIS Elements software and all downstream analyses were performed on these projections. In experiments involving intensity measurements (GFP11 signal accumulation, IFIT1 transcription and IRF3-BFP nuclear translocation) analyzed channels were corrected for photobleaching using the ‘bleach correction’ plugin in ImageJ.

### Data analysis

#### smFISH analysis

To calculate the fraction of infected cells that was positive for antiviral gene expression, the number of infected cells was first determined based on the EMCV FISH signal. A cell was considered infected if specific EMCV FISH signal was detected. However, only if a cell had >50 EMCV smFISH spots it was considered infected, because at 16 h.p.i., considerable release of viral particles from infected cells results in significant smFISH signal originating from virus particles on the outside of a cell. We note that beyond ∼8 h.p.i. the amount of viral genomes in infected cells was frequently too high to allow detection of individual viral RNAs. Cells with high EMCV signal for which the number of genomes could no longer be counted were also scored as EMCV positive. In the experiment with dipyridamole treatment (Fig. 1E) no cells with >50 vRNAs could be detected and instead all cells in the large image were evaluated for IFIT1 and IFNB1 expression (Again, viral particles on the outside of the cell preclude accurate quantification of the number of vRNAs in the cell by smFISH). The number of IFIT1 and/or IFNB1 spots was determined for each infected cell. The number of spots was determined using the ‘Spot Counter’ plugin in ImageJ. Detection settings were optimized for each experiment and for each channel individually due to experimental variation in smFISH labeling, and automated analysis was manually curated for each measurement. For IFIT1, IFNB1 and Puro-P2A-SNAP smFISH, spots that were unusually bright (>2,5 fold mean intensity of single spots) and non-spherical were not scored as individual mRNAs, as such foci likely originated from dye aggregates. At approximately 200 spots per cell, considerable overlap in spots in the maximum intensity projection impairs accurate spot detection and therefore 200 spots per cell was set as an upper limit for quantification.

If cells were partially outside the field of view or if cells were blurred at an image stitch during large image construction, they were not included in the analysis.

### Viral load measurement

The viral load was determined using the cytosolic fluorescence intensity of EMCV-Atto488 FISH staining. The mean intensities of 10-20 ROIs of 20×20 pixels, that were randomly positioned in the cytoplasm without overlap, were determined and the average of the measurements was calculated. The number of ROI measurements was chosen so that >80% of the cell’s cytoplasm was ultimately part of an intensity measurement. For each repeat an average cellular background signal intensity was determined in the same manner using uninfected cells. Average background signal was subtracted from average cellular EMCV signal intensities. In order to compare viral load between different experiments these values were normalized to the average intensity of the ten cells with the highest signal in the experiment and this normalized value was multiplied by 1000.

### smFISH spot intensity

To determine the distribution of IFIT1 and IFNB1 smFISH spot intensities, the intensity of all spots (max. 30/cell) in a random region of a IFIT1/IFNB1+ cell was determined. For this a 4×4 pixel ROI centered around the middle of a spot was used to measure the mean fluorescence intensity. Mean intensity of an adjacent 4×4 pixel ROI without spot was determined and subtract from the spot intensity. Non-spherical/overlapping spots and putative transcription sites (based on quantity, nuclear localization and higher intensity) were excluded from the analysis. Background subtracted spot intensities were normalized to the average spot intensity in the cell.

For transcription site (TS) intensity measurements based on IFIT1 and Puro-P2A-SNAP smFISH probes, the intensity of 5 individual (24xPBS) IFIT1 mRNAs was determined in the same manner as described above for IFIT1 and IFNB1 mRNAs. The TS intensity was subsequently determined by measuring the mean intensity in a 7×7 pixel ROI positioned over the center of the TS and subtracting the mean intensity of a similarly sized ROI positioned directly adjacent to the TS (in a region devoid of smFISH spots). TS intensity was divided by the average intensity of the 5 individual mRNAs to calculate the relative TS intensity. A spot was considered a TS if it was localized in the nucleus and had a relative intensity of >2,5fold based on either IFIT1 or Puro-P2A-SNAP labeling.

### VIRIM quantification

Annotation of viral infection phases based on SunTag spots in VIRIM was performed as described previously (Boersma, et al., 2020). In brief, GFP spots are considered viral translation sites based on their size, mobility and intensity. For example, when cells express both GFP-STAb and GFP(1-10) a fraction of cells present with large cytosolic GFP spots that are not viral translation sites (they are present in uninfected cells as well). However, these spots can be readily discriminated from viral translation sites based on their larger size and slower mobility.

The start of VIRIM phase 3 is defined as the moment where one or more viral translation site(s), as visualized by SunTag labeling, re-appear after being absent during VIRIM phase 2 (initial replication phase). Because of the imaging interval of 5 or 10 min and the relatively short duration of phase 3 (∼30 min), a steep increase in the number of SunTag spots between frames is typically observed during VIRIM phase 3. This steep increase in the number of SunTag spots was used to pinpoint the start of phase 3 in those cells in which accurate calling of VIRIM phase 1 (and 2) was challenging.

### VIRIM in combination with smFISH

Various parameters were determined from the VIRIM live imaging and smFISH staining. Start of VIRIM phase 3, number of IFIT1 and IFNB1 spots and EMCV viral load was determined as described above.

Only cells for which complete VIRIM history and successful smFISH staining was available were analyzed. Cells for which time lapse imaging data could not be faithfully linked smFISH data (for instance because high cell density) were excluded from analysis. Cells that underwent mitosis were excluded from the analysis if mitosis took place just before the start of VIRIM phase 3 (<30 min after completion of cytokinesis) or if mitosis occurred between start of phase 3 and the end of the movie. For EMCV(L^WT^) infected cells, only positions in which an IFIT1+ cell was present were analyzed.

### 24xPBS IFIT1 (and CMV) transcription site intensity measurements

Identification of 24xPBS IFIT1 transcription sites during live imaging was based on the following criteria: 1) an IFIT1 TS is a spherical (diffraction limited) spot and considerably smaller in size compared to typical nucleoli (which are also enriched in mCherry signal – see for instance Fig. 4A). An IFIT1 TS fits within a 5×5 pixel ROI. 2) IFIT1 TSs show slow and highly confined diffusion within the nucleus. 3) an IFIT1 TS emerges during the course of infection, i.e. they are absent at the start of infection and not present in uninfected cells. 4) The mCherry fluorescence intensity of a IFIT1 TS fluctuates over time, in contrast to mCherry signal originating from nucleoli and aggregates. 5) An IFIT TS is present for a prolonged period (>60min, in a minimum of 6 frames).

Multinucleated cells or cells that formed syncytia during the movie were excluded from analysis.

The time point of the first appearance of an mCherry spot that meets these criteria is considered as the onset of IFIT1 transcription.

The intensity of IFIT1 TSs is determined by measuring the mean intensity of a 5×5pixel ROI positioned over the center of the TS and subtracting the mean intensity of an ROI of the same size positioned directly adjacent to the TS from this value. If a TS overlaps (partially) with a nucleoli then background subtraction is performed by measuring the intensity of an ROI in the direct vicinity of the TS that has a comparable fraction of nucleolar overlap. From the intensity time trace the area under the curve was determined using the trapezoidal rule: for this, the average intensity value between consecutive time points was determined and multiplied by the time interval. These values were summed to determine the AUC of an IFIT1 TS intensity time trace. Cells with AUC > 5000 a.u. were considered positive for IFIT1 transcription. To quantify IFIT1 transcriptional activity at the onset of transcription, the average IFIT1 TS intensity in the first hour after the appearance of an IFIT1 TS was determined. If a TS was temporarily absent during this 1h time period, an intensity value of zero was included in the average calculation.

Intensity of the 24xPBS CMV TS intensity was determined in a similar fashion to IFIT1 TS intensity measurements. Compared to IFIT1, CMV TS identification differed in one aspect: CMV TSs are present at the start of the movie and remain present throughout the movie. To be included in the analysis at least 66% of the frames must have a detectable CMV TS. Average CMV intensity traces were smoothened by applying a moving average with a window size of 5 time points.

### Split-GFP intensity measurements

To measure GFP signal originating from split-GFP reconstitution, the mean cytosolic GFP intensity was measured at every time point. For this a 25×25 pixel ROI was positioned in the perinuclear region of the cell at 3 non-overlapping positions and the average of these measurements was calculated. If fluorescent aggregates were present in the cell (caused by aggregation of the STAb-GFP and GFP1-10, see also “VIRIM quantification”), these were avoided. To subtract baseline GFP signal originating from the GFP-STAb, the average cytosolic GFP intensity in the first 2h of the movie was subtracted from all values. In some cases morphological changes to the cell occurred during the movie that strongly affected GFP intensity measurements, for example during cell death or detachment at the end of infection. In such cases, measurements after the morphological changes occurred were excluded from further analysis.

### Automated split-GFP measurement and cluster analysis

Nuclear segmentation was performed based on the nuclear signal of PCP-NLS-mCherry using cellpose (Stringer, et al., 2021). The mean GFP pixel intensity for each nuclear mask at each time point was computed. Single cells were tracked over time using the btrack algorithm (Ulicna et al., 2021). Segmentation and tracking results were displayed in napari and the performance of the algorithms was manually curated for a subset of positions. A track length of at least 40 time points was chosen as a quality threshold, resulting in a total number of 1430 tracks from 3 independent experiments.

Single cell split-GFP tracks were smoothed by applying a moving average with a window size of 10 time points. The smoothed single cell traces were then clustered using the dynamic time warping algorithm from the dtw R package (Sardá-Espinosa, 2019). The resulting distance matrix was used for hierarchical clustering using average linkage and split into eight clusters. Clusters 4-8 were removed because they contained only a small number of traces (7, 5, 4, 2 and 1, respectively), resulting in three main clusters corresponding to the non-infected traces (flat shape curve, n = 440) and two sigmoidal curve shapes, differing in their growth rate and plateau height (split-GFP Low/Medium n= 842, split-GFP High n=129).

### IRF3-BFP nuclear translocation

To determine the intensity ratio of nuclear/cytosolic IRF3-BFP, BFP signal intensity was measured in the nucleus and cytosol in the same manner as cytosolic split-GFP signal intensity was determined (i.e. the average of 3 mean intensity measurements using a 25×25 pixel ROI was determined; see section “Split-GFP intensity measurements”). Background correction was performed by determining the mean intensity of a 25×25 pixel ROI positioned at a place in the FoV where no cells were located and this value was subtracted from all nuclear and cytosolic intensities. The nucleocytosolic IRF3-BFP ratio was determined for every time point and time traces were aligned to the start of IFIT1 transcription (or start of phase 3 in the case of Fig. 6C). Ratio time traces were normalized to the average nucleocytosolic ratio between 2 to 7 h before the start of IFIT1 transcription (or start phase 3).

To quantify the translocation efficiency, the AUC of the nucleocytosolic ratio time traces before normalization was determined using the trapezoidal rule starting from 6h after start of phase 3 until the end of the movie (this time point was chosen, because at 6h after the start of phase 3 the first infected cells display IRF3-BFP nuclear translocation). The AUC was divided by the number of frames that were included in the AUC calculation to correct for different trace durations.

IRF3-BFP nuclear translocation was defined using the following requirements: 1) an increase in the nucleocytosolic ratio is observed in multiple, consecutive frames (spanning >30 min, in a minimum of 3 frames). 2) during at least two frames, the increase in the nucleocytosolic ratio was at least 0.1 unit. The moment of translocation was defined as the first time point of a series of frames that fulfill these requirements. If no IRF3-BFP nuclear translocation was observed according to these criteria, cells were not included in the analysis to determine the timing difference between the moment of translocation and start of IFIT1 transcription (Fig. 6D).

### Statistical analysis

Statistical tests were performed making use of a p-value of 0.05 as a cutoff for significance. The type of test and the type of error bars used in figures are indicated in the figure legends.

To extract descriptive parameters from the split-GFP intensity time traces (Fig. S4), a logistic growth curve was fitted on the (average) split-GFP traces using the following general equation:

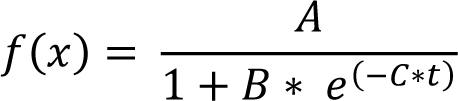

Where A represents the plateau value and C is the logistic growth rate. The maximum slope was calculated from the plateau value and C parameter using the following equation:

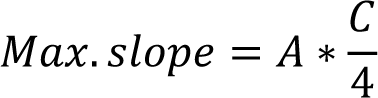

The mean squared error (MSE) was calculated to determine the quality of fit.

(Linear) regression analysis was performed in the GraphPad PRISM software. To quantify the extent of correlation in Fig. 1F and S2C the Pearson’s correlation coefficient (r) was determined. In Fig. 3 linear regression was performed excluding observations where the number of mRNAs was above the detection limit (>200 mRNAs). The 95% confidence interval of the linear fit and the coefficient of determination (R2) was determined to assess the quality of the linear regression.

Supplemental data with Fig. 1 A) Fraction of 5xSunTag-EMCV(L^WT^) or 5xSunTag-EMCV(L^Zn^) infected cells expressing 20 or more IFIT1 mRNAs at 8 h.p.i. (n =3 independent experiments) * indicates p < 0.05 using independent samples T-test. B) Fraction of EMCV(L^Zn^) or 5xSunTag-EMCV(L^Zn^) infected cells expressing >20 IFIT1 or >10 IFNB1 mRNAs at 16 h.p.i. (n = 577 and 532 cells for EMCV(L^Zn^) and 5xSunTag EMCV(L^Zn^) respectively in 4 independent experiments) n.s. indicates non significant, p > 0.05 using independent samples T-test. C) Histogram of the number of IFIT1 and IFNB1 smFISH spots in cells which were not incubated with virus (n = 144 cells in 3 independent experiments) D) Fraction of cells that express more than 20 IFIT1 mRNAs in cells that were treated with different concentrations of IFN for 24 h in the presence or absence of the the JAK1/3 inhibitor Tofacitinib (TOFA.) (n = 3 independent experiments). E) IFIT1 smFISH spot intensity distribution in IFNB1- and IFNB1+ cells (n = 600 and 570 spots respectively from 3 independent experiments) Bars and error bars indicate average ± s.e.m. in all panels

Supplemental data with Fig. 2 A) Histogram of the time between the moment of virus addition to the cell culture medium and start of phase 3 (n = 399 infections in 3 independent experiments). B) Relative IFIT1 (black, left y-axis) and IFNB1 (red, right y-axis) mRNA levels in HeLa cells expressing STAb-GFP infected with 5xSunTag-EMCV(L^Zn^) at different time points as determined by qPCR. mRNA levels are expressed relative to expression at the moment of virus addition (t = 0) (n = 3 experiments). C) Viral genome abundance in HeLa cells expressing STAb-GFP infected with 5xSunTag-EMCV(L^Zn^) at different time points as determined by qPCR (n = 3 experiments). D) Average viral load of 5xSunTag-EMCV(L^Zn^) infected IFIT1-/IFNB1-, IFIT1+/IFNB1-, and IFIT1+/IFNB1+ cells at different time periods since the start of phase 3 (n = 243, 122, and 34 cells respectively in 3 independent experiments) * indicates p < 0.05 using a two-way ANOVA test, n.s. indicates non significant. E) Scatter plot showing the number IFIT1 (left) and IFNB1 (right) mRNAs and viral load at 16 h.p.i. (r indicates Pearson’s correlation coefficient, n = 399 cells in 3 independent experiments). Error bars indicate s.e.m. in all panels

Supplemental data with Fig. 3 A) Schematic representation of the 24xPBS reporter knock in in the IFIT1 locus and genotyping results. To assess correct integration of the reporter cassette, a 5’ and 3’ PCR fragment was amplified from genomic DNA using a primer that binds to the IFIT1 genomic sequence and a primer that binds to the reporter sequence. PCR reactions were analyzed by gel electrophoresis and PCR fragments of correct size (indicated with red asterisk) were purified and subjected to sanger sequencing to confirm correct integration (right panel). To determine the number and intensity of IFIT1 transcription sites, both on the edited and unedited allele, 24xPBS-IFIT1 k.i. cells were infected with 5xSunTag-EMCV(L^Zn^) for 16 h and subjected to smFISH using probes targeting the IFIT1 coding sequence and a probe set targeting the PURO-P2A-SNAP region of the reporter mRNA. B) Shows a histogram of the number of untagged (black bars) or PP7-tagged (red bars) IFIT1 transcription sites (n = 196 cells in 4 independent experiments).

Supplemental data with Fig. 4 A) Maximum slope values of the split-GFP intensity time trace of IFIT1-(black dots) and IFIT1+ cells (red dots) in individual repeats of the experiment shown in Fig. 4C. Dots connected by a line represent values obtained in a single independent experiment. See method section for explanation about the quantification of max. slope. * indicates p < 0.05 using paired samples T-test. B) Maximum slope values of split-GFP time traces from all individual infections in IFIT1- and IFIT1+ cells. See method section for explanation about the quantification of max. slope. Red bars indicate average and s.d. (n = 46 IFIT1+ and 117 IFIT1-infections in 4 independent experiments). * indicates p < 0.05 using independent samples T-test. C) Characteristics of GFP medium/low and GFP high infections. n indicates the total number of infections that were assigned to either group in 3 independent experiments. Fraction of infections indicates the relative proportion of either type of infection. MSE = mean squared error

Supplemental data with Fig. 5 24xPBS-IFIT1 k.i. cells stably expressing STAb-GFP, PCP-mCherry-NLS, and a CMV-driven 24xPBS reporter RNA were infected with 5xSunTag-EMCV(L^Zn^) and imaged for 16 h Average CMV transcription site intensities during 5xSunTag-EMCV(L^Zn^) infection in IFIT1+ (red line) and IFIT1-(black line) cells aligned to the start of phase 3 (n = 24 and 24 IFIT1+ and IFIT1-cells respectively in 3 independent experiments) Shaded areas indicate s.e.m.

Supplemental data with Fig. 6 A) Schematic representation of the BFP knock in in the IRF3 locus and genotyping results. To assess correct integration of the BFP coding sequence, a 5’ and 3’ PCR fragment was amplified from genomic DNA using a primer that binds to the IRF3 genomic sequence and a primer that binds to the BFP sequence. PCR reactions were analyzed by gel electrophoresis and PCR fragments of correct size (indicated with red asterisk) were purified and subjected to sanger sequencing to confirm correct integration (right panel). B) Bar graph of the fraction of EMCV(L^Zn^) infected IRF3-BFP k.i. or parental HeLa cells expressing >20 IFIT1 mRNAs. IRF3-BFP k.i. and HeLa cells were infected with 5xSunTag-EMCV(L^Zn^) and subjected to smFISH for IFIT1 mRNA at 16 h.p.i. Bars and error bars indicate average ± s.e.m. (n = 426 and 399 infected cells for HeLa and IRF3-BFP respectively in 3 independent experiments) n.s. indicates p > 0.05 using independent samples T-test.

Video S1 Movie of live-cell virus imaging using VIRIM combined with smFISH for IFIT1, IFNB1 and EMCV. HeLa cells expressing STAb-GFP were infected with 5xSunTag-EMCV(L^Zn^) and imaged for 16 h. After that, cells were fixed and smFISH staining was performed for IFNB1 mRNA (magenta), IFIT1 mRNA (cyan), and positive strand EMCV vRNA (EMCV(+), white). For the VIRIM imaging, a maximal intensity projection of 9 Z-slices was generated. Time is indicated in hours:minutes since the start of image acquisition. Images of the smFISH staining were projected on top of the last frame of the VIRIM imaging.

Video S2 Movie of live-cell virus imaging using VIRIM and split-GFP in combination with IFIT1 transcription imaging. 24xPBS-IFIT1 k.i. cells expressing STAb-GFP, GFP(1-10) and PCP-mCherry-NLS were infected with GFP11-5xSunTag-EMCV(L^Zn^) and imaged for 16 h. Left panel (green) shows VIRIM and split-GFP signal, middle panel (magenta) shows IFIT1 transcription signal (first appearance of IFIT1 TS at 11h10) and the right panel shows the merge of both imaging channels. For both channels, 9 Z-slices were acquired and processed to generate a maximal intensity projection. Time is indicated in hours:minutes since the start of image acquisition. Images from this movie are also shown in the example images in Fig. 4A.

Video S3 Movie of live-cell imaging of IFIT1 transcription and IRF3-BFP nuclear translocation. IRF3-BFP and 24xPBS-IFIT1 k.i. cells stably expressing STAb-GFP and PCP-mCherry-NLS were infected with 5xSunTag-EMCV(L^Zn^) and imaged for 16 h. Left panel (in grey) shows IFIT1 transcription signal (first appearance of IFIT1 TS at 13h00) and the right panel (displayed using the “inferno” lookup table) shows the IRF3-BFP signal. For the IFIT1 transcription imaging, 9 Z-slices were acquired every 10 min. and processed to generate a maximal intensity projection. For IRF3-BFP, a single Z-slice was acquired every 60 min. Time is indicated in hours:minutes since the start of image acquisition. Images from this movie are also shown in the example images in Fig. 6A.

