## Supplemental Data for "Heterogeneity in viral replication dynamics shapes the antiviral response"

### SUPPLEMENTAL DATA WITH FIG. 1

**S1A**

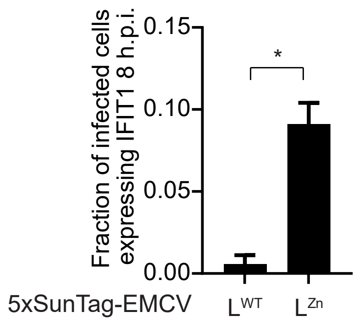


**S1B**

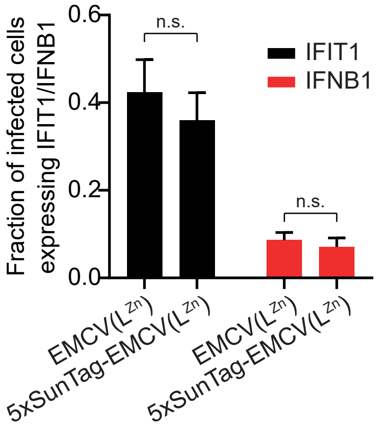


**S1C**

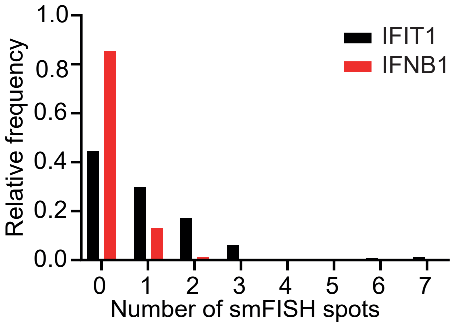


**S1D**

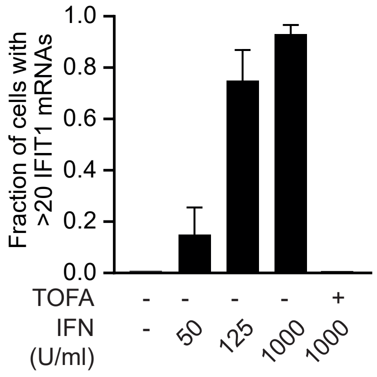


**S1E**

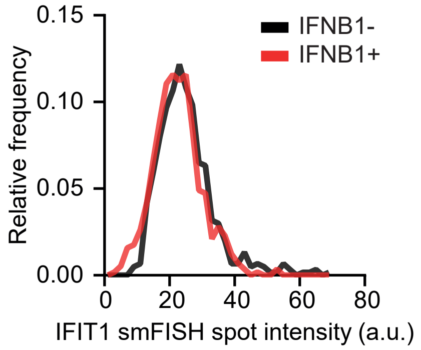

### SUPPLEMENTAL DATA WITH FIG. 2

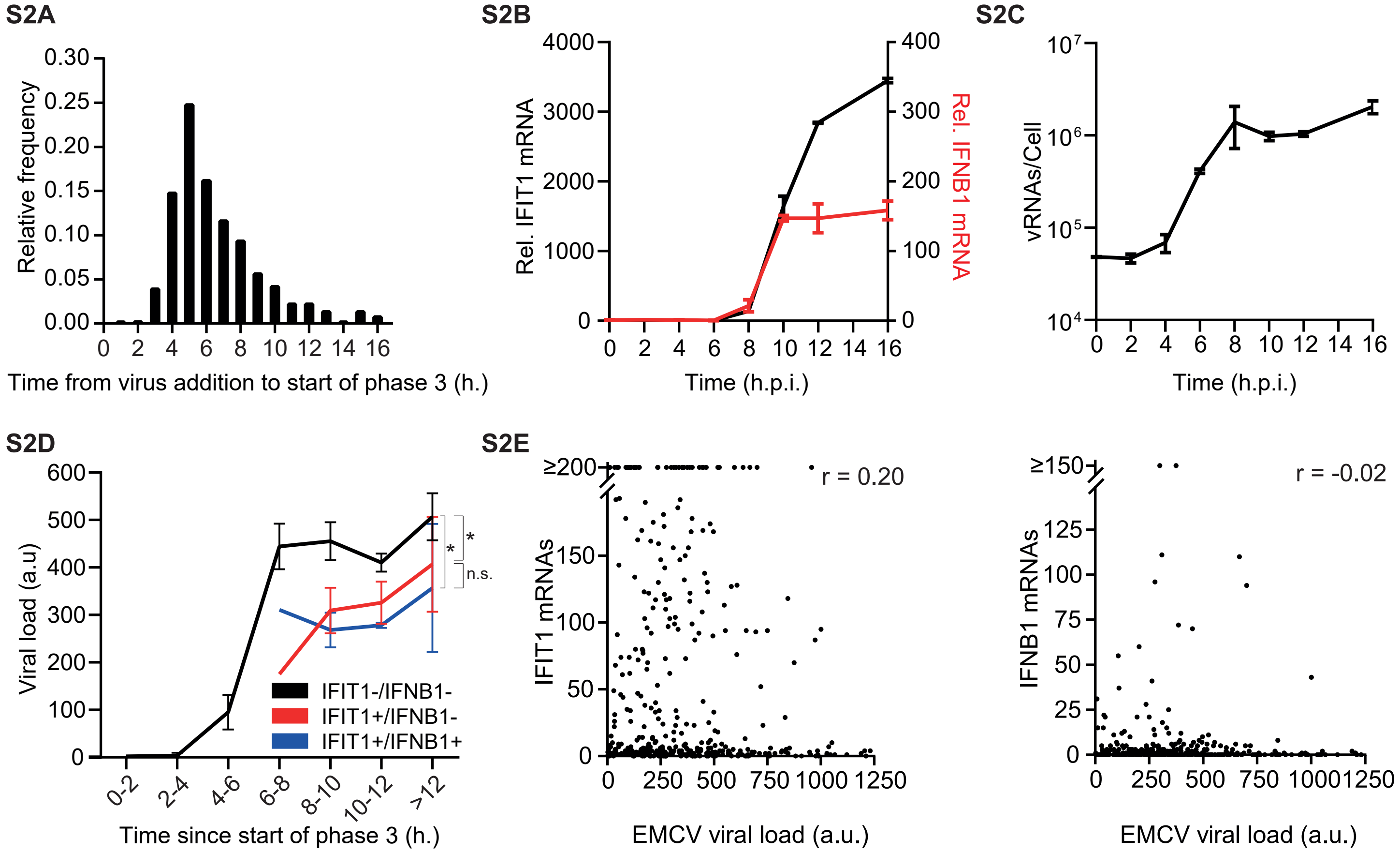

### SUPPLEMENTAL DATA WITH FIG. 3

### S3A

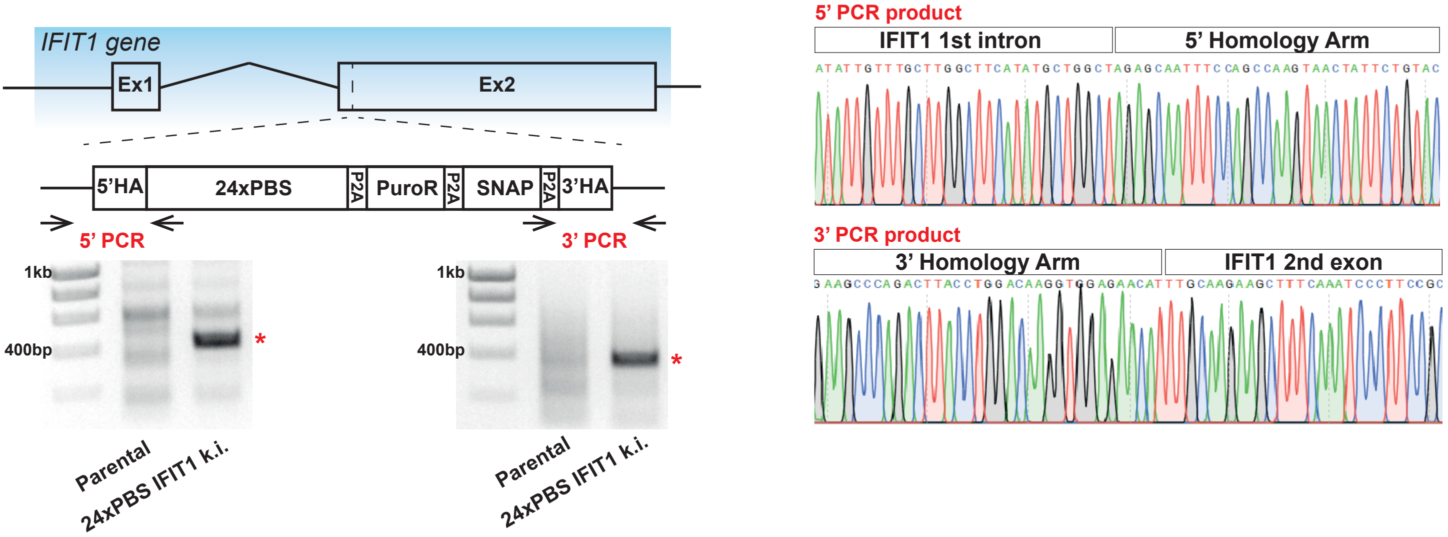


### S3B

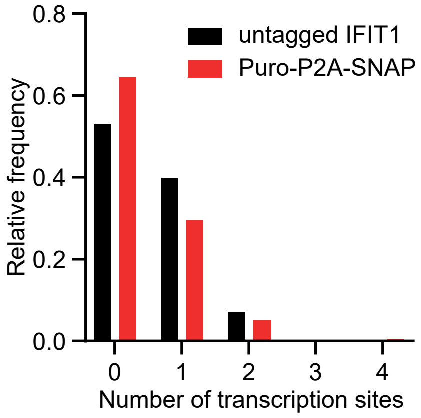

**SUPPLEMENTAL DATA WITH FIG. 4**

**S4A**

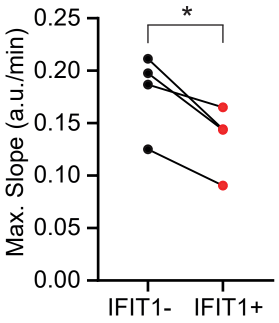


**S4B**

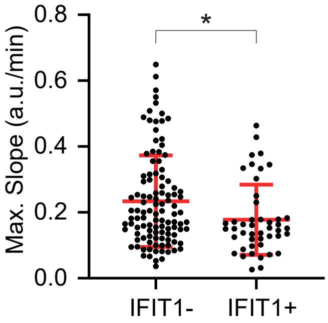


**S4C**

| | split-GFP | |
|---|---|---|
| | Low/Medium | High |
| n (infections) | 842 | 129 |
| Fraction of infections | 0.87 | 0.13 |
| Logistic Fit parameters | | |
| Maximum slope | 0.083 | 0.173 |
| Plateau value | 17.97 | 41.23 |
| MSE | 0.82 | 1.30 |

**SUPPLEMENTAL DATA WITH FIG. 5**

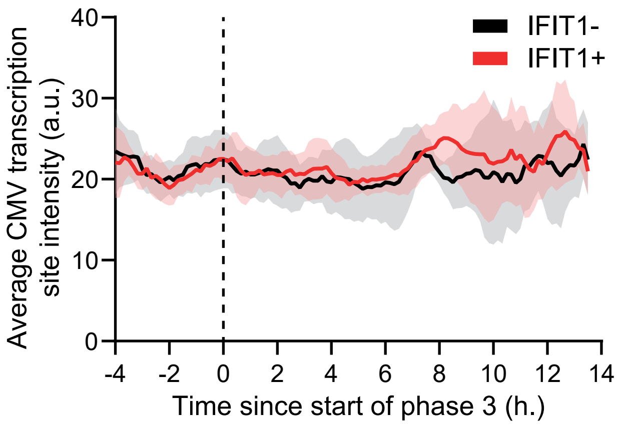

### SUPPLEMENTAL DATA WITH FIG. 6

### S6A

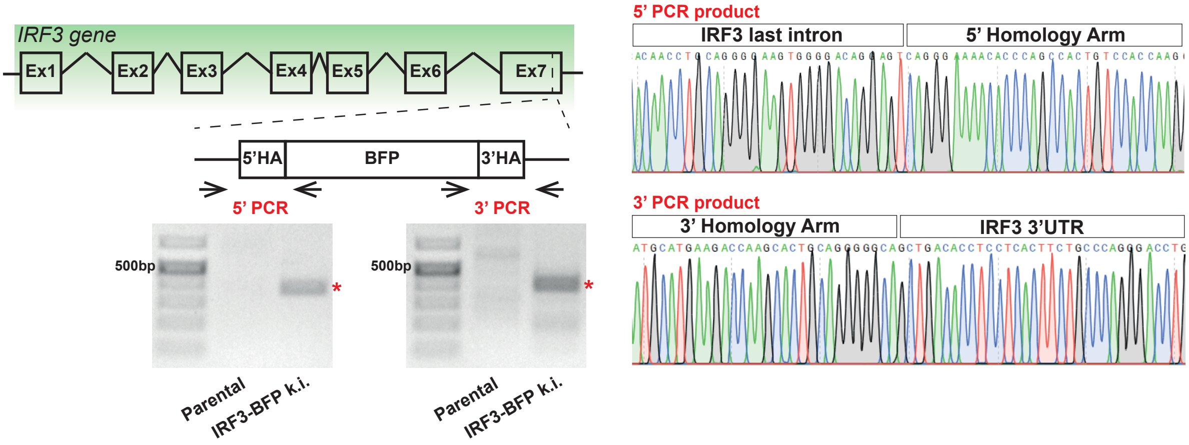


### S6B

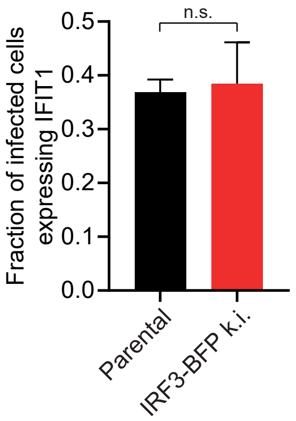
